# Differential substrate specificity of ERK, JNK, and p38 MAP kinases toward Connexin 43

**DOI:** 10.1101/2023.12.30.573692

**Authors:** Lauren P. Latchford, Liz S. Perez, Jason E. Conage-Pough, Reem Turk, Marissa A. Cusimano, Victoria I. Vargas, Sonal Arora, Sophia R. Shienvold, Ryan R. Kulp, Hailey M. Belverio, Forest M. White, Anastasia F. Thévenin

## Abstract

Phosphorylation of connexin 43 (Cx43) is an important regulatory mechanism of gap junction (GJ) function. Cx43 is modified by several kinases on over 15 sites within its ∼140 amino acid-long C-terminus (CT). Phosphorylation of Cx43CT on S255, S262, S279, and S282 by ERK has been widely documented in several cell lines, by many investigators. Phosphorylation of these sites by JNK and p38, on the other hand, is not well-established. Indeed, ERK is a kinase activated by growth factors and is upregulated in diseases, such as cancer. JNK and p38, however, have a largely tumor-suppressive function due to their stress-activated and apoptotic role. We investigated substrate specificity of all three MAPKs toward Cx43CT, first by using purified proteins, and then in two cell lines (MDCK - non-cancerous, epithelial cells and porcine PAECs – pulmonary artery endothelial cells). Cx43 phosphorylation was monitored through gel-shift assays on an SDS-PAGE, immunodetection with phospho-Cx43 antibodies, and LC-MS/MS phosphoproteomic analyses. Our results demonstrate that p38 and JNK specificity differ from each other and from ERK. JNK has a strong preference for S255, S262, and S279, while p38 readily phosphorylates S262, S279, and S282. While we confirmed that ERK can phosphorylate all four serines (255, 262, 279, and 282), we also identified T290 as a novel ERK phosphorylation site. In addition, we assessed Cx43 GJ function upon activation or inhibition of each MAPK in PAECs. This work underscores the importance of delineating the effects of ERK, JNK, and p38 signaling on Cx43 and GJ function.

## INTRODUCTION

Gap Junctions (GJs) are cell structures that uniquely allow for direct cell-cell communication within tissues, and it has been well established that GJ intercellular communication (GJIC) is crucial for a multitude of cellular outcomes; GJs regulate the coordination of development, cellular homeostasis, and tissue function.^1–4^ In vertebrates, GJs are made up of integral membrane proteins termed connexins (Cx). Each Cx molecule spans the membrane bilayer four times, with both the N and the C-terminus in the cytoplasm. Six connexin monomers oligomerize into a half GJ termed a hemichannel while trafficking through the secretory pathway. When they arrive at the plasma membrane, they interact with the hemichannels found in the neighboring cell’s plasma membrane, forming a complete GJ channel.^5,6^

Connexin oligomerization, forward trafficking, GJ channel formation, and opening, as well as GJ closing, internalization, and degradation, are precisely regulated through posttranslational modifications and interactions with many binding partners (protein chaperones, scaffolds, ubiquitination, and endocytosis machinery).^1,7,8^ The protein life cycle of Cx43 (the most ubiquitously expressed connexin), is regulated through phosphorylation/dephosphorylation events on over 15 sites by at least 9 kinases within the flexible and largely unstructured C-terminal tail. This includes mitogen-activated protein kinases (MAPKs), Src, protein kinases A and C (PKA and PKC), Akt, casein kinase 1 (CK1), and cyclin-dependent kinase 1 (CDK1/cdc2).^8^

MAPK family consists of three distinct signaling pathways: extracellular signal-regulated kinase (ERK), p38, and c-Jun N-terminal kinase (JNK).^9^ Four MAPK phosphorylation sites were previously identified within the C-terminal region of Cx43: S255, S262, S279, and S282. There is a large body of evidence of ERK phosphorylation at these four sites in multiple cell lines.^8,10–19^ While some studies of Cx43 phosphorylation by JNK and p38 have been conducted, none focused on the identification of specific serine residues.^20–23^ Even though these parallel MAPK pathways are highly conserved, they respond to distinct stimuli, leading to diverse, and often cell line-specific outcomes. ERK signaling is activated mostly through growth factor receptors, resulting in changes in cell proliferation and migration,^24^ while JNK and p38 pathways respond to stress, such as osmotic shock, hypoxia, and reactive oxygen species.^9,25^

We systematically evaluated the differences in substrate specificity of each MAPK toward Cx43. First, we treated purified GST-Cx43CT mutants with purified and active MAPKs through kinase assays with purified protein components. We measured Cx43 phosphorylation through three complementary techniques: SDS-PAGE gel mobility shifts, immunoblotting with phosphoCx43-specific antibodies, and liquid chromatography with tandem mass spectrometry (LC-MS/MS) phosphoproteomics. Importantly, we utilized phosphomimetic and phospho-dead mutants at MAPK sites as positive and negative controls, respectively, taking advantage of Cx43’s ability to undergo electrophoretic mobility shifts, even when using phosphomimetic mutations (S to E). Our findings map out the phosphorylation preferences of each MAPK toward S255, S262, S279, and S282 on Cx43, indicating that each MAPK has a differential specificity. Importantly, we identified T290 as a novel ERK phosphorylation site on Cx43. Moreover, we assessed Cx43 phosphorylation by MAPKs in two cell lines: MDCKs and PAECs (epithelial and endothelial cell lines, respectively). Our results demonstrate that ERK phosphorylation of Cx43 predominates in MDCKs, while JNK and p38 are the primary MAPKs that phosphorylate Cx43 in PAECs. Finally, we demonstrate that Cx43 GJs close when ERK, JNK, or p38 are activated, while inhibition of MAPKs leads to an increase in cell-cell communication.

## RESULTS

MAPKs are proline-driven kinases and recognize the following phosphorylation consensus motif in their substrates: P-X-**S/T**-P, with the phospho-acceptor serine or threonine at the [0] position (**Fig. 1A**).^1,26–28^ Two sites (S279 and S282) in both the human and rodent Cx43 sequence fully conform to this phosphorylation motif, while S255 lacks the proline in the [-2] position but does have the proline in the [+1] position. In addition, S255 conforms to a cyclin-dependent kinase motif ^29^ (S-P-X-R/K, **Fig. 1B**) and has been observed to be phosphorylated by a cyclin-dependent kinase cdc2 at the onset of mitosis.^30,31^ More recently, S255 phosphorylation by MST1 (mammalian STE20-like kinase) was detected in endothelial cells during atherosclerosis^32^ - a surprising finding given MST’s strong preference against S/T residues that are followed by a proline in the +1 position.^33^ Unlike S255, S279, and S282 sites, serine 262 in the human sequence of Cx43 is not a conserved site due to lack of proline at both [-2] and [+1] positions in the human sequence (**Fig. 1B**). However, it is not surprising that S262 phosphorylation has been observed by many researchers because either a rodent cell line was used or a rodent (typically, rat) sequence of Cx43 was expressed in human cells or studied *in vitro*.^12,34^ Nonetheless, there is plenty of evidence that MAPKs can phosphorylate substrates that do not fully conform to the P-X-S/T-P motif, also relying on key docking site sequences that may be available both on the substrate and on the MAPK itself.^26–28,35,36^ It is therefore important to systematically assess each MAPK’s ability to phosphorylate these serine sites on Cx43 **(Fig. 1C)** rather than relying on the presence of canonical phosphorylation motifs **(Fig. 1A and B)**.

**Figure 1:**
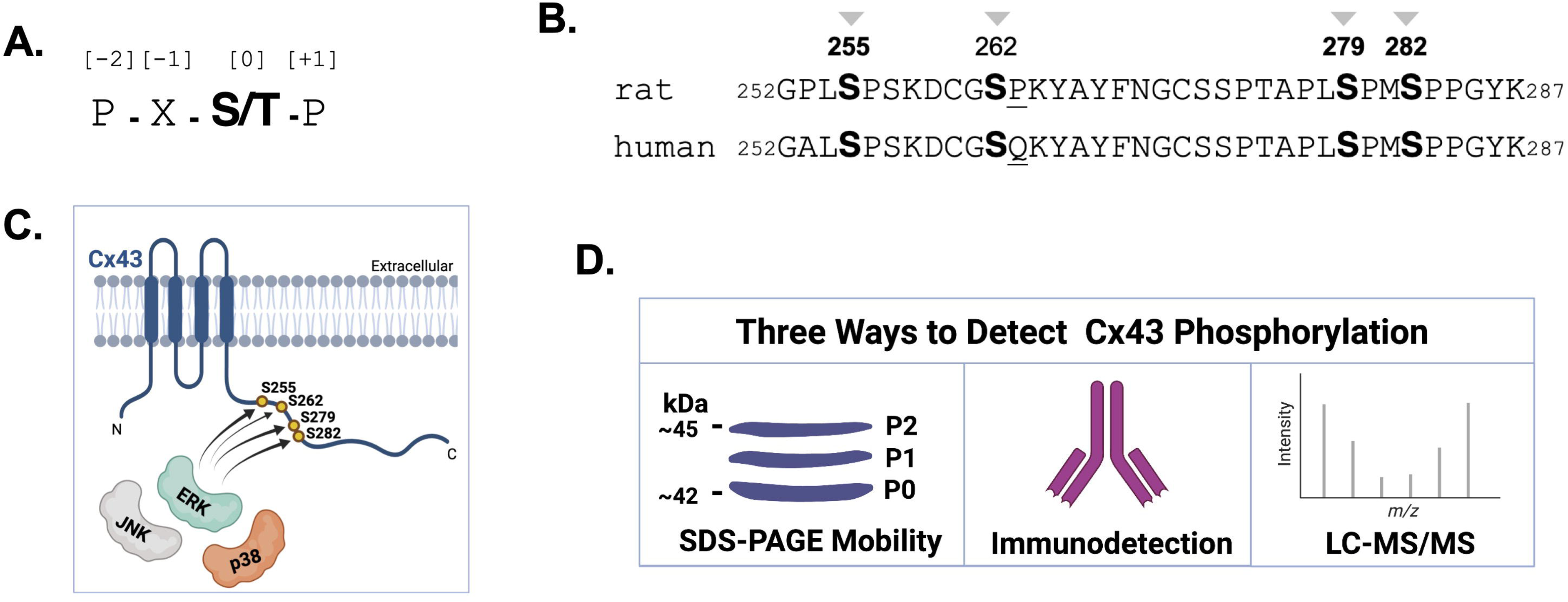
**A.** Phosphorylation motif for MAPK substrates, where S/T is the phosphoacceptor residue at position 0, and there is a preference for a proline residue at +1 and -2 positions. X – any amino acid. **B.** Human vs. rat amino acid sequence alignment within the Cx43CT that harbors the relevant serine residues (shown in bold). While S279 and S282 fully conform to the MAPK motif in both human and rat connexin 43 sequences, S255 in the human sequence lacks a proline at the -2 position in humans. S262, while followed by a proline at +1 in the rat Cx43, it lacks a proline in the human sequence (underlined). Cx43 used in this study is from a rat gene unless otherwise specified. Sequence alignment was generated using Clustal Omega Multiple Sequence Alignment tool. **C.** Overall question of this study: Do ERK, JNK, and p38 MAPKs display differential phosphorylation preference toward serine residues (255, 262, 279, and 282) on the Cx43 CT? **D.** Cx43 phosphorylation was monitored by three techniques: electrophoretic mobility shifts on an SDS-PAGE from an unphosphorylated, P0 form to the phosphorylated P1 and P2 forms (see text for detail), immunodetection with phosphospecific Cx43 antibodies (to identify Cx43 phosphorylated at S255, S262, S279, or S282), and through use of liquid chromatography and tandem mass spectrometry (LC-MS/MS) of trypsin-digested Cx43 protein samples. Figures C and D were generated in BioRender.

We chose to assess Cx43 phosphorylation with three complementary approaches: detection with phosphospecific antibodies, electrophoretic mobility shifts, and liquid chromatography, tandem mass spectrometry (LC-MS/MS) analyses **(Fig. 1D)**. Phosphospecific antibodies against serine residues on Cx43 have been successfully developed, commercialized, and used by many researchers.^10–12,37^ In addition, it has been well-established that Cx43 undergoes electrophoretic mobility shifts when phosphorylated at specific serine residues. At least three electrophoretic forms have been observed: P0, P1, and P2, where unphosphorylated Cx43 migrates in the P0 isoform, with phosphorylation by some kinases shifting to P1, P2, (even to P3 in some cell lines) or remaining at P0.^1,7,23,38^ These shifts have also been recapitulated by replacing serine sites with phosphomimetic mutations. For example, a triple phosphomimetic mutant (S255D/S279D/S282D) of recombinantly expressed and affinity purified Cx43 C-terminus demonstrates a shift to the P1 form, while single and double S255D/S262D remain in P0.^39^ Thus, shifts to slower-migrating forms (when combined with the use of phospho-specific antibodies and serine site mutants) represent a valuable readout of Cx43 phosphorylation (**Fig. 1D**). Despite the utility of phosphospecific antibodies, users are limited to observing only the phosphorylation sites recognized by a given antibody. Moreover, phosphoantibody performance can vary, with occasional non-specific binding to the unphosphorylated form or other proteins in the mixture. Given these shortcomings, we sought to complement gel shifts and phosphoantibody usage with mass spectrometry-based phosphoproteomics - an unbiased discovery method.

### Kinase Assays Between Pure MAPKs and GST-Cx43CT Proteins Indicate Differences in Phosphorylation Preference

Recombinant ERK2, p38⍺, and JNK2⍺2 were purified from *E.coli* in their active form by coexpression with their upstream activators. Histidine-tagged ERK was coexpressed with a constitutively active MKK1. Robust ERK activation was seen in samples that contained both ERK and MKK1, while active ERK antibodies detected significantly lower levels of active ERK when MKK1 was absent. Total ERK expression levels were also detected with histidine tag antibodies and were highest in IPTG-treated samples (**Fig. S1A**). Milligram amounts of ERK were purified from the soluble fraction through nickel affinity chromatography and ERK retained its activity throughout the purification process (**Fig. S1B**). Active, GST-tagged p38⍺ was produced through coexpression with its upstream activator, MKK6. The highest levels of active GST-tagged p38 were seen in the soluble protein fraction **(Fig. S1C**). GST-tagged, active p38⍺ was purified by glutathione-affinity chromatography (**Fig. S1D**). Active, histidine tagged JNK2⍺2 was produced by expression with its two upstream activators: a constitutively active form of MEKK (MEKKC) and MKK4. Robust expression of active JNK was seen in the soluble *E.coli* fraction induced with IPTG (**Fig. S1E**). Histidine-tagged, active JNK was of high purity and retained its activity throughout the purification process (**Fig. S1F**).

To conduct kinase assays between each pure MAPK and Cx43, we expressed GST-tagged Cx43 C-terminus in *E.coli* (WT, phospho-dead and phospho-mimetic single, double and triple mutants: S255A, S255E, S279A/S282A, S279E/S282E, S255A/S279A/S282A, S255E/S279E/S282E) and purified them using glutathione affinity chromatography (**Fig. S1G**) to be used as substrates in phosphorylation reactions with active ERK, JNK, and p38 (**Fig. 2A**).

**Figure 2:**
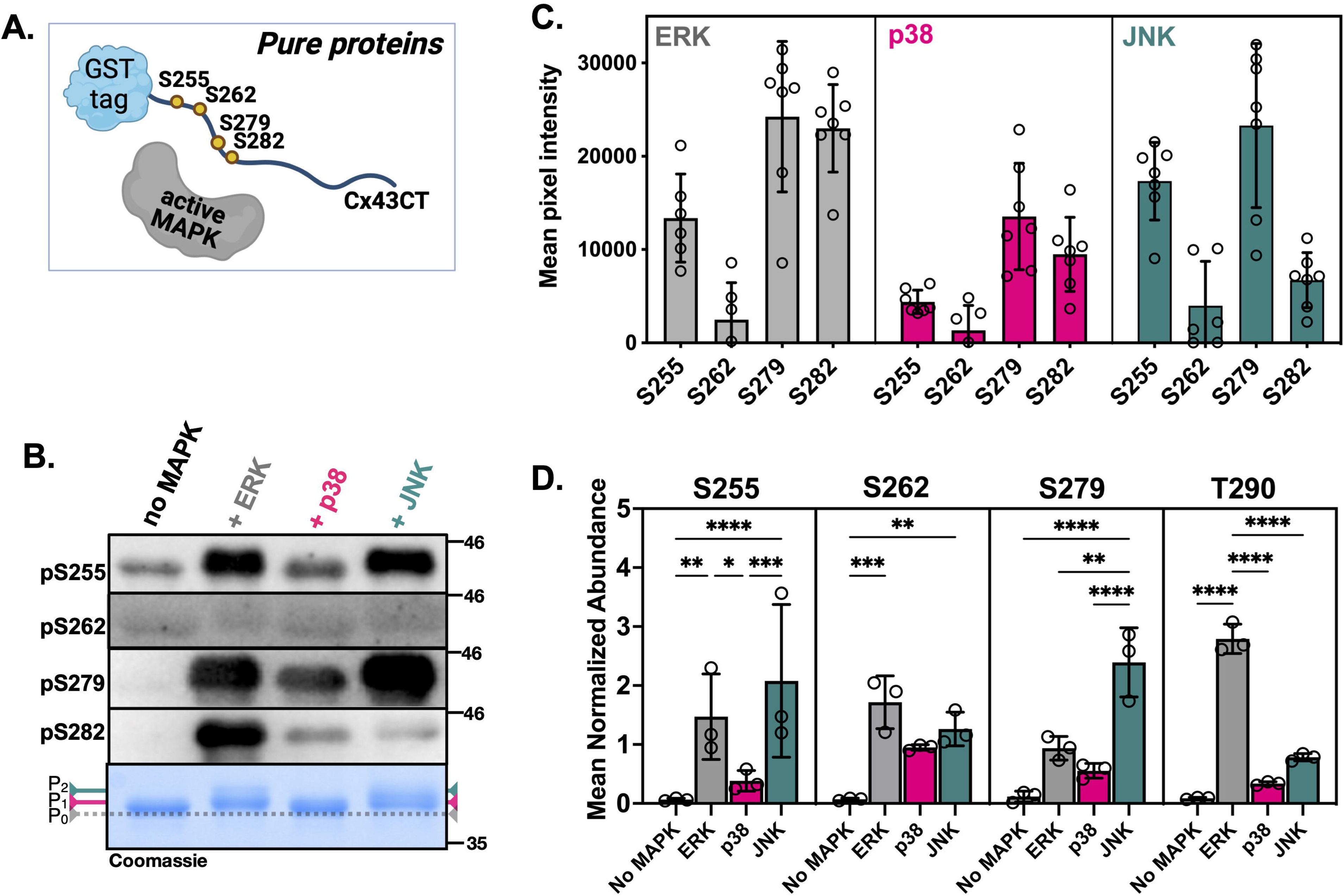
**A.** Our approach to studying Cx43 phosphorylation by MAPKs through use of recombinantly-produced and purified GST-tagged Cx43CT protein (6 µM) treated with purified and active MAPKs (His- ERK2, GST-p38ɑ, and His-JNK2 at 0.5 µM each). **B.** WB analyses with phosphoserine-specific Cx43 antibodies on GST-Cx43CT (WT) kinase assay samples with each indicated active MAPK Total Cx43 levels and mobility shifts were assessed with Coomassie staining. The reaction without active kinase is included as a negative control. A representative example is shown. **C.** Quantification of mean pixel intensities in ImageJ of phosphoserine Cx43 signal from kinase assays in (A). Data represent seven independent replicates (n=7), and error bars are shown in SD (standard deviation). The antibody background signal from the ‘no MAPK’ control was subtracted from each phosphoserine site signal. Because different phosphoserine antibodies were used, statistical significance was not assessed. Total Cx43 in the Coomassie stain in B was used as a loading control. **D.** Phosphoproteomics was used to measure the phosphorylation of GST-Cx43CT after incubation with active forms of the indicated MAPKs. Bars represent the relative abundance of phosphopeptides containing the indicated Cx43 phosphorylation sites. Values are normalized to the average abundance measured for GST-Cx43 across each sample condition. Data represent three replicates (n=3) per condition and statistically significant differences were determined by two-way ANOVA Tukey multiple comparisons test: p < 0.05(*), p < 0.005 (**), p < 0.0005 (***), and p < 0.0001 (****). Error bars are shown in SD (standard deviation). Figure A was generated in BioRender.

GST-tagged WT Cx43CT was initially tested with each pure kinase to identify kinase assay conditions that could be used across all three MAPKs (**Fig. S2A-C**). We incubated 6µM WT GST- Cx43CT, triple A, or triple E mutants in a time-dependent (0-60 minutes, **Fig. S2A and S2B**) and in ERK enzyme concentration-dependent (0-1µM) manner - data not shown. It was evident that a full electrophoretic shift was observed within 30 min. We also tested time-dependent phosphorylation of WT GST-Cx43CT with active JNK (0-60min) at 0.5 µM JNK, establishing that JNK-treated Cx43CT was no longer shifting after 15-30 minutes (**Fig. S2B)**. Unlike JNK and ERK-treated GST-Cx43CT, p38-treated GST-Cx43CT failed to demonstrate any changes in electrophoretic mobility (across 0-1µM kinase concentration range, over the course of 1 hour) - **(Fig. S2C)**. Given the abovementioned observations, for subsequent experiments, we chose the following kinase reaction conditions across all three MAPKs and every Cx43CT mutant: 60 min incubation of 0.5 µM kinase and 6 µM GST-Cx43CT.

Next, we conducted Western blotting (WB) analyses of Cx43 phosphorylation with phospho-specific antibodies against S255, S262, S279, and S282 (**Fig. 2A-C**). Antibodies against pS255 are not commercially available and were shared by Dr. Paul Lampe’s laboratory (Fred Hutch Cancer Center, Seattle, WA). Antibodies against pS279 and pS282 were highly specific, while pS255 antibodies displayed some level of non-specific detection in the control (no MAPK) sample **(Fig. 2B)**. pS262 antibodies were the most non-specific, with all four samples (including the negative control) displaying similar levels of signal across multiple repeats. We attempted further dilutions of pS262 antibodies to 1:2000 and 1:5000 but were unable to achieve better specificity (data not shown). It is important to note that in these experiments, the highest antibody specificity was achieved when using a less sensitive chemiluminescence reagent (see Methods).

Clear differences in phosphorylation of GST-Cx43CT WT by each MAPK were observed (**Fig. 2B**, quantitated in **2C**). ERK phosphorylated all three serine sites (255, 279, and 282), while p38 had lower overall levels of Cx43 phosphorylation, toward all sites, and particularly toward S255. JNK, on the other hand, had robust levels of S255 and S279 phosphorylation, with lower levels of S282 phosphorylation. However, we felt that statistical analyses of these experiments would not be appropriate given the likely differences in the specificity of each Cx43 phospho- serine antibody.

In addition to phospho-specific antibody experiments, we processed kinase assay samples for analysis using phosphoproteomics. This approach allowed us to strengthen evidence for the phosphorylation sites detected by western blotting, while also potentially identifying Cx43 phosphorylation sites that lack phospho-specific antibodies. Additionally, our use of isobaric mass tags in these experiments facilitated relative quantitation of the amount of phosphorylation of a given site across multiple conditions. We were able to detect several (>15) singly and multi-phosphorylated C-terminal Cx43 peptides using this method (**Table S1**). To increase our confidence in phosphosite assignment, limited validation of mass spectra was performed for the phosphorylation sites characterized with phosphoantibodies **(Figs. S3-S6**). The mass spectrometry results largely mirrored the antibody studies, with JNK and ERK demonstrating substantially more activity for Cx43 at each phosphorylation site than p38 (**Fig. 2D, Table S1**). Together, phospho-specific antibody and mass spectrometry data suggest similar JNK preference for S255 compared to ERK. Phosphoproteomics also provided a clear advantage over the use of the S262 antibody, indicating that is phosphorylated by all three MAPKs, with a slight preference for ERK phosphorylation at this site. While both approaches captured the preference of JNK for the S279 phosphorylation site, mass spectrometry data showed this phosphorylation event was strongly associated with JNK activity. Due to the high signal intensity of the singly phosphorylated S279 Cx43 peptide, we were unable to validate and confidently identify peptides with single S282 phosphorylation or double S279/S282 phosphorylation. However, raw data point to potential preferential ERK phosphorylation at S282 (**Table S1**). Strikingly, our discovery mass spec approach identified a strongly ERK-driven phosphorylation event at T290 (**Fig. 2D**, **Table S1**, **Fig. S6**). This site is particularly intriguing since it does not adhere to the canonical P-X-S/T-P or S/T-P motifs. Overall, the phosphoproteomic data corroborate the phosphoantibody findings while providing a more quantitative assessment of differences among the MAPK conditions.

To better compare the effects of each MAPK on the electrophoretic mobility of Cx43, we treated 6µM WT GST-Cx43CT with pure and active ERK, p38, or JNK (0.5µM each) **(Fig. 3A)**. Quantification of gel shifts by ImageJ (**Fig. 3B-C**, n=7) indicated that while ERK and JNK-treated GST-Cx43CT proteins undergo a shift to P1 and P2 forms, p38-treated Cx43 does not display a significant shift as compared to the negative control (**Fig. 3C**) or as compared to GST-Cx43CT WT treated with active ERK (**Fig. S2D**).

**Figure 3:**
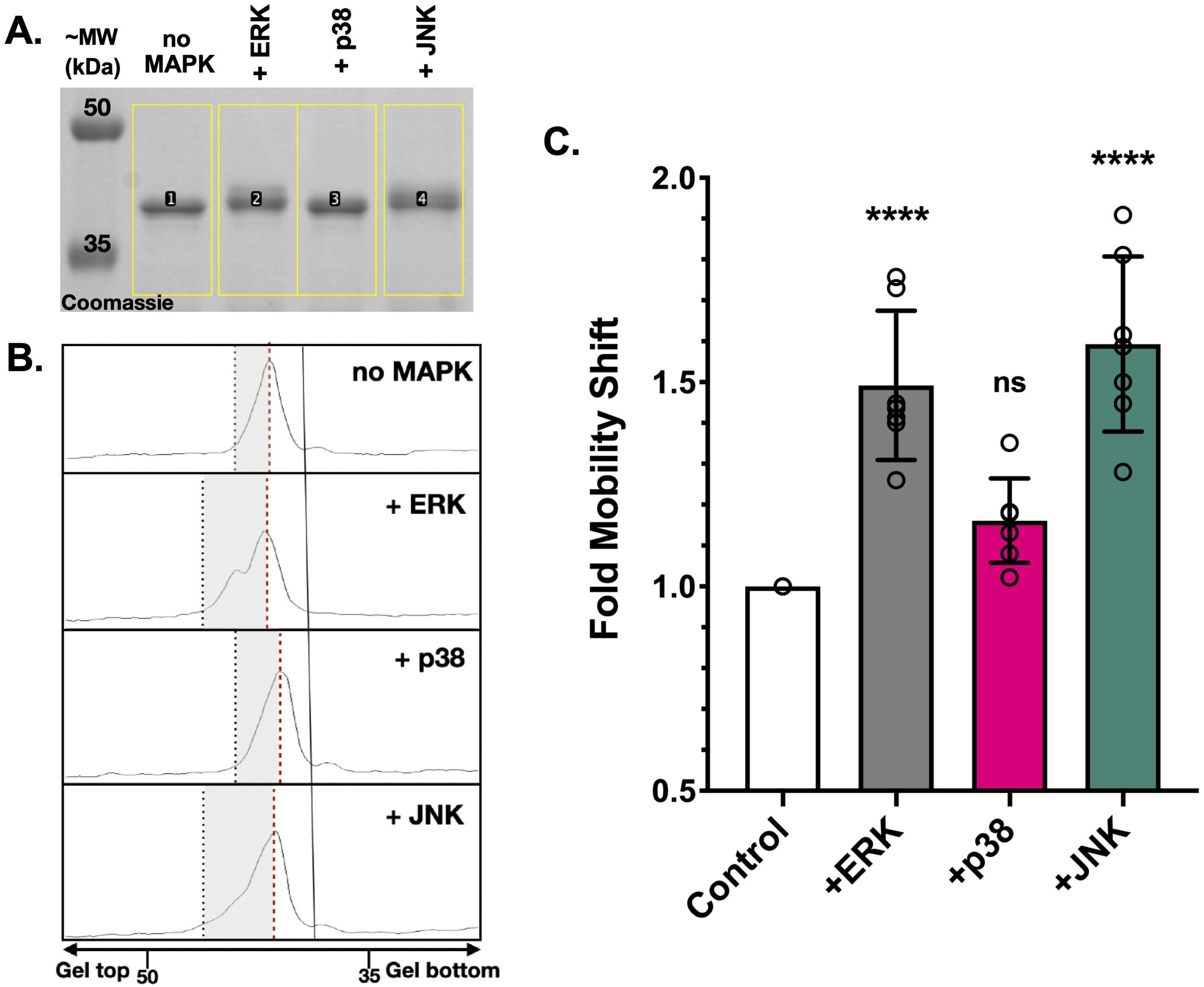
**A.** A representative SDS-PAGE image of kinase reactions between GST-Cx43CT (WT, 6 µM) and each active MAPK (0.5 µM). A reaction sample lacking active MAPK is included for comparison. Yellow boxes and lane numbers indicate areas selected in the ImageJ software to visualize and quantify Cx43 mobility shifts. **B.** Representative image of pixel intensity peaks obtained from (A) in ImageJ. Peak maxima are indicated with a dashed red line, while the left-hand side of each peak (from the maximum to the minimum) are highlighted in gray rectangles. The width of each gray rectangle (indicative of a change in Cx43CT mobility) was measured in Keynote. **C.** Statistical analyses of GST-Cx43CT mobility shifts. Data represent seven biological replicates (n=7) per kinase reaction and statistically significant differences were determined by ordinary one-way ANOVA: p < 0.0001 (****), ns-not significant. Error bars are shown in SD (standard deviation).

To better identify the effects of each Cx43 phosphorylation event on electrophoretic mobility changes from P0 to P1 and P2 isoforms, we utilized a larger series of our purified phosphomimetic and phospho-dead mutants at S255, 279/282, and T290 (**Fig. S1G)**. Given ERK’s ability to phosphorylate all five sites (S255, S262, S279, S282, and T290 – **Fig. 2D**), we chose this MAPK for the kinase assays with the GST-Cx43CT mutants (**Fig. 4A and 4B**). Kinase assays of each Cx43CT protein with or without the addition of active ERK were analyzed by WB analyses against each serine site (255, 262, 279, and 282), as well as by electrophoretic mobility shifts using total Cx43 antibodies (**Fig. 4A-B**, summarized in **4C**). Total Cx43 western blot indicated that WT, 255A, 279A/282A, and triple A mutants were in the P0 form without the addition of active ERK (**Fig. 4A**). S255E and double E mutant (no ERK) were observed in the P1 form, while the triple E mutant (no ERK) was in the P2 form. Upon treatment with active ERK, WT Cx43CT exhibited a smear (likely representative of a combination of all three forms (P0, P1, and P2). 255A shifted to P1, while 255E shifted to P2. ERK treatment of S279A/S282A led to both P0 and P1 forms, while the double E mutant exhibited both P1 and P2 forms. Triple A mutant treated with active ERK remained unchanged and stayed in the P0 form, while the triple E mutant stayed at P2. Taken together, these results indicate that a shift to P2 requires phosphorylation at all three sites (255, 279, and 282), and in the absence of S255 phosphorylation or 279 and 282 phosphorylation, Cx43 only shifts Cx43 to the P1 form (**Fig. 4A** and summarized in **4C**).

**Figure 4:**
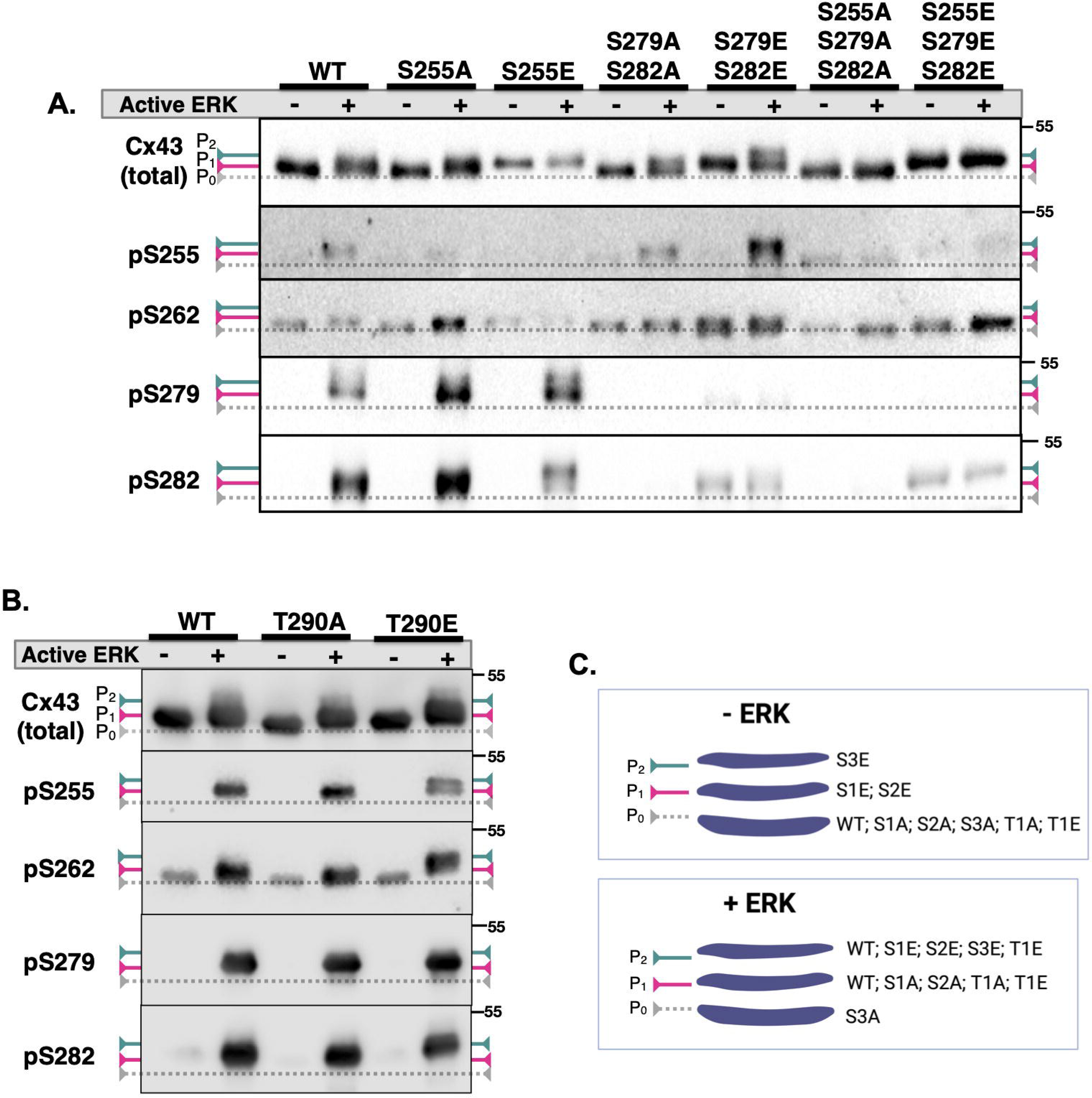
Electrophoretic mobility shift and WB analyses of kinase assays between active ERK and pure GST-Cx43CT. Positions of the unphosphorylated (P0) and phosphorylated (P1 and P2) are indicated. **A.** Assays with serine mutants. **B.** assays with T290 mutants. The same reaction conditions were used in A and B as in **Figs. 2-3**). **C.** Summary of results from (A) and (B) under two conditions: without and with active ERK. When active ERK was not included, unphosphorylated WT and all serine-to-alanine mutants (S1A, S2A, S3A) were detected in the P0 form (S1A, S2A, S3A). S255E (S1E) and S279E/S282E (S2E) were detected in the P1 form, and the triple E mutant (S3E) was in the P2 form. In addition, both T290A and T290E mutants (T1A and T1E) ran at the P0 form. Once treated with active ERK, the S3A mutant remained at P0, S1A and S2A shifted to P1, while S1E and S2E shifted to P2. The S3E mutant remained in the P2 form, similar to the ‘no ERK’ conditions. The T1A mutant shifted to P1, while the T1E mutant was seen both in P1 and p2. Figure C was generated in BioRender.

Assessment of serine phosphorylation through immunodetection with phospho-specific Cx43 antibodies indicated that S255 is phosphorylated in the WT, 279/282 double A, and double E mutants, demonstrating a very specific signal from these antibodies. Moreover, the pS255 band was observed in the P2 form, concurring with the results of the gel shifts in the total Cx43 WB (**Fig. 4A**). Interestingly, levels of pS255 were higher in the double E mutant than in the double A mutant, even though total Cx43 levels were similar. A fair level of nonspecific signal for S262 phosphorylation was observed in the ‘no ERK’ samples for all the mutants and the WT. These results yet again point to the low quality of this antibody. However, a large increase - well above the background signal - was observed in the triple E mutant treated with active ERK as compared to that mutant without ERK. This result demonstrates that S262 phosphorylation by ERK is highly efficient once the other three sites have been phosphorylated (**Fig. 4A**). Moreover, an increase in pS262 signal is seen in S255A mutant treated with ERK. However, this may partially be due to higher levels of total Cx43 in that sample (see total Cx43 WB). Overall, pS262 phosphorylation signal is observed in the P1 form across all mutants.

Robust levels of S279 phosphorylation were observed in the WT and in both S255 mutants. The WT treated with active ERK exhibited a smear (likely indicative of a shift to both P1 and P2). S255A mutant displayed S279 phosphorylation in the P1 form, while S255E mutant contained both P1 and P2 forms. Indeed, these results reinforce the observation from the total Cx43 signal that S255 phosphorylation is necessary for a shift to P2 when S279 is phosphorylated (**Fig. 4**).

Lastly, the detection of S282 phosphorylation indicated that while WT and S255A displayed a shift to P1, the S255E mutant had a shift to P2, yet again demonstrating that the combination of S255 and S282 provides a shift to P2. Given the observation that P2 bands in S255E mutant are seen with both S279 and S282 antibodies, it is likely that all three phosphorylation are needed to shift Cx43 to P2. The double and the triple-A mutants did not display any S282 signal, while some level of the nonspecific signal was observed in both the double and triple E mutants, regardless of the presence of active ERK.

To test whether the phosphorylation status of T290 would have any effect on ERK phosphorylation of the serine sites (255, 262, 279, 282), we generated T290A and T290E mutants of our GST-Cx43CT (**Fig. S1G**) and utilized them in kinase assays with active ERK (**Fig. 4B**). We observed an increase in the electrophoretic mobility of the T290E mutant when probed with total Cx43, pS255, pS262, and pS282 antibodies. However, mimicking phosphorylation at T290 (T290E vs. T290A) did not cause an electrophoretic shift in the absence of ERK (**Fig. 4B**, summarized in **4C**).

### Activation and Inhibition of Individual MAPKs in Mammalian Cells

To test Cx43 phosphorylation by MAPKs expressed in two mammalian cell lines, we first identified cellular conditions under which we could activate a single MAPK at a time (**Fig. 5**). We chose to use Madin Darby Canine Kidney cells (MDCK), a cell line that does not express any endogenous Cx43 but can be transfected with full-length WT Cx43 and phosphomimetic/phospho-dead mutants.^40^ In addition, we also assessed levels of endogenous Cx43 phosphorylation in pPAECs (porcine Pulmonary Artery Endothelial Cells). In MDCKs, ERK was activated by cell treatments with the epidermal growth factor (EGF), while in PAECs, we compared ERK activation upon EGF vs. VEGF (vascular endothelial growth factor) treatments. To activate p38 and JNK signaling pathways, both cell lines were treated with anisomycin^41^ (**Fig. 5**). Unfortunately, both JNK and p38 are involved in cellular responses to stress and we were not able to identify activators of only the JNK or p38 pathway.^20,22,23^ Thus, we utilized inhibitors of p38 (SB202190) and JNK (SP600125) in an attempt to downregulate the kinase activity of each of these MAPKs. We utilized MAPKAPK-2/MK2 (mitogen-activated protein kinase-activated protein kinase 2) phosphorylation as a readout of p38 activity and c-Jun phosphorylation to measure JNK activity.^25,42,43^ In addition, it has been observed that even though SB202190 and SP600125 are competitive inhibitors that prevent downstream phosphorylation of p38 and JNK substrates, respectively, there is evidence that binding of these inhibitors will also affect levels of p38 and JNK phosphorylation by their upstream MAP2Ks (MEK3/6 and MEK4/7, respectively).^44–47^ Thus, we also monitored levels of p38 and JNK phosphorylation as partial readouts of these two pathways. To inhibit ERK activity, we utilized a dual MEK1/MEK2 inhibitor (U0126)^44^ and measured levels of phosphorylated ERK as a readout of this signaling pathway (**Fig. 5**).

**Figure 5:**
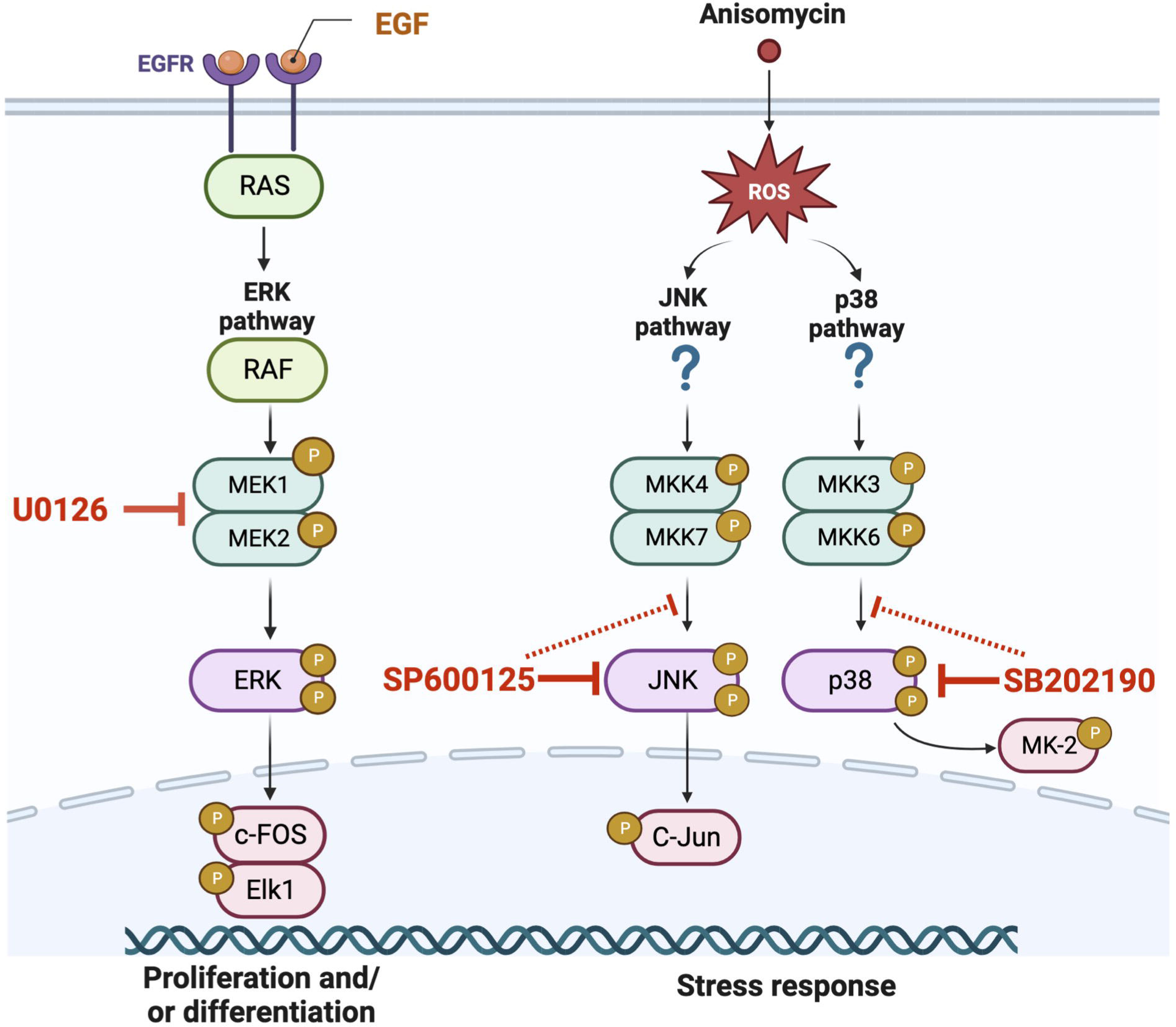
MAPK signaling pathways, their activators, and inhibitors relevant to this study. ERK signaling pathway is activated through the action of the EGF receptor (EGFR), Ras GTPase, MAP3K, Raf and MAP2Ks, MEK1 and MEK2. To inhibit ERK in cells, an inhibitor of MEK1 and MEK2 was used (U0126). Anisomycin was utilized as an activator for both JNK and p38 stress pathways, through activation of MEK4/7 and MEK3/6, respectively. To inhibit JNK or p38, SP600125 and SB202190 were used, respectively. These inhibitors are known to target JNK and p38 directly, preventing phosphorylation of downstream targets. However, there is a smaller effect on JNK and p38 phosphorylation by their respective MEKs. Changes in JNK and p38 activity were tested by monitoring the phosphorylation of their targets (c-Jun and MK2) and levels of phosphorylated JNK and p38. Figure was adapted from an existing template in BioRender.

We were successful at establishing activation and inhibition conditions for each MAPK in MDCKs (**Fig. 6A**), as measured through levels of phosphorylated ERK, cJun, and MK2. Importantly, total levels of each MAPK, c-Jun, and MK-2 remained largely unchanged across all treatment conditions (**Figs. 6B**). ERK activation was robust upon treatments of starved cells with EGF, while treatment with the MEK inhibitor drastically diminished levels of active ERK (quantitated in **Fig. 6C**). Basal levels of active ERK were also detected when cells were grown in full media (FBS – **Fig. 6C**). As judged by the levels of phosphorylated c-Jun, JNK was the most active upon treatment with Anisomycin, and JNK activity decreased when cells were treated with JNK inhibitor alone, or together with the p38 inhibitor (quantitated in **Fig. 6D**). Activation of p38 (as judged by the levels of phosphorylated MK-2) was the most robust upon treatment with anisomycin and diminished when cells were treated with p38 inhibitor alone or together with the JNK inhibitor (**Fig. 6E**).

**Figure 6:**
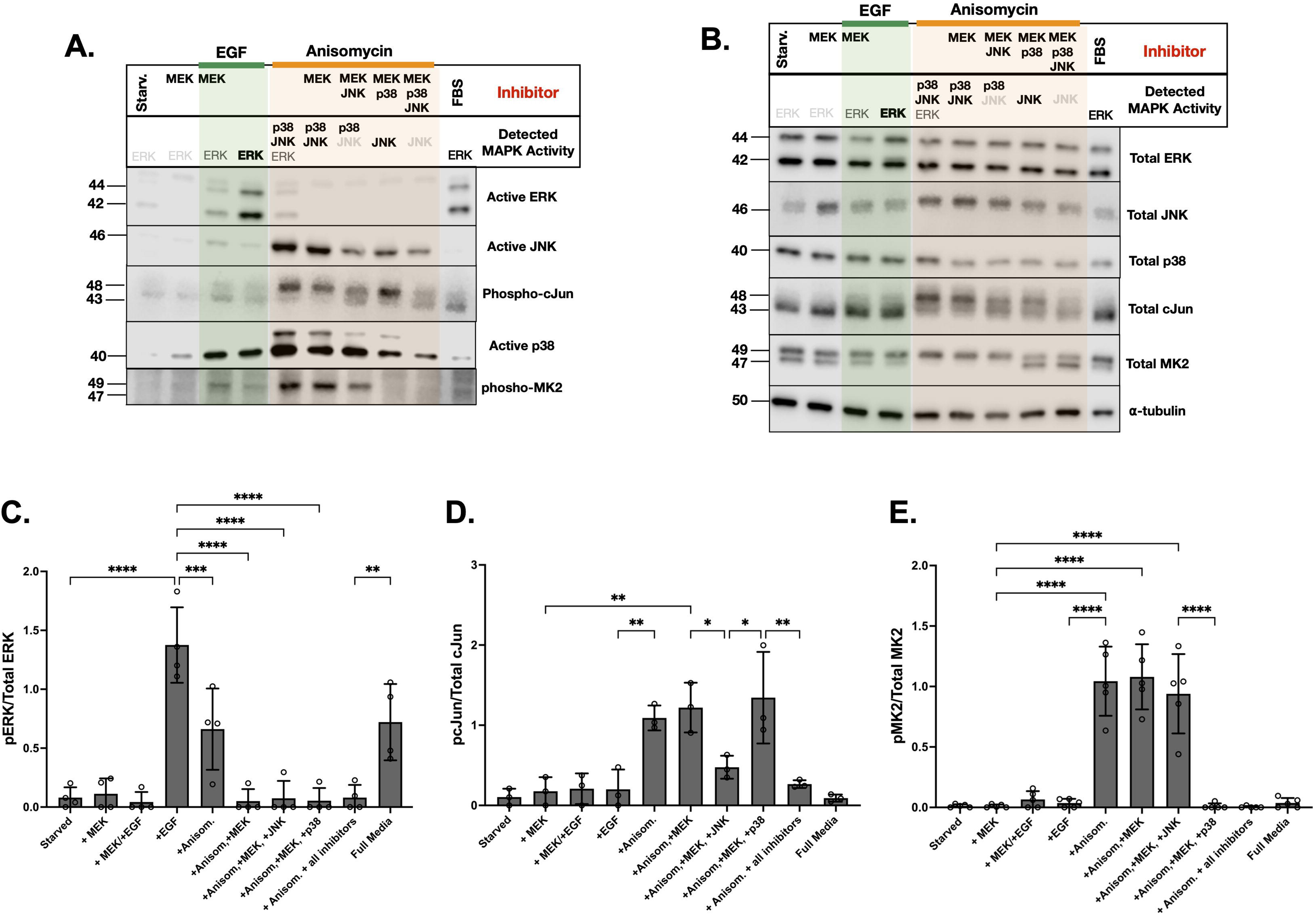
Assessment of MAPK activities in MDCK lysates. **A.** One representative treatment repeat: Western blots of active ERK, JNK, p38, and phosphorylated downstream targets of JNK (c-Jun) and p38 (MK2) in MDCK lysates treated with EGF (green highlight), Anisomycin (orange highlight), and various combinations of kinase inhibitors (as listed at the top). Detected MAPK activity for each condition is summarized using darker, bolded font (higher activity) vs. lighter, thinner font to indicate lower/basal activity. **B.** Control Western blots of total MAPKs, c-Jun, MK-2, and the α-tubulin loading control in MDCK lysates. The same samples and treatment conditions were used as in A. **C.** Quantification of active ERK vs total ERK levels from western blots in A and B. **D.** Quantification of JNK activity through detection of phosphorylated vs. total cJun blots from A and B. **E.** Quantification of p38 activity through detection of phosphorylated vs. total MK-2 from A and B. Statistically significant differences in C-E were determined by ordinary one-way ANOVA: p < 0.05(*), p < 0.005 (**), p < 0.0005 (***), and p < 0.0001 (****). Error bars are shown in SD (standard deviation), from at least three independent cell treatments.

We were also successful at establishing activation and inhibition conditions for each MAPK in PAECs (**Fig. 7A**) as measured through levels of phosphorylated ERK, cJun, and MK2. Yet again, total levels of each MAPK, c-Jun, and MK-2 remained largely unchanged across all treatment conditions (**Fig. 7B**). To activate ERK PAECs, we utilized both EGF and VEGF treatments (**Fig. 7A**). All other treatments were identical to what we report in MDCKs (**Fig. 6**). The levels of ERK activation were statistically similar for both growth factors, even though PAECs are an endothelial cell line. High active ERK signal was also observed in cells grown in full media (FBS) (**Fig. 7C**). Highest levels of active JNK and p38 (as judged by phosphorylated c-Jun and MK-2 signals, respectively) were seen upon anisomycin treatment, while activity of each kinase dropped when cells were treated with the respective kinase inhibitors (**Figs. 7D** and **E**, respectively).

**Figure 7:**
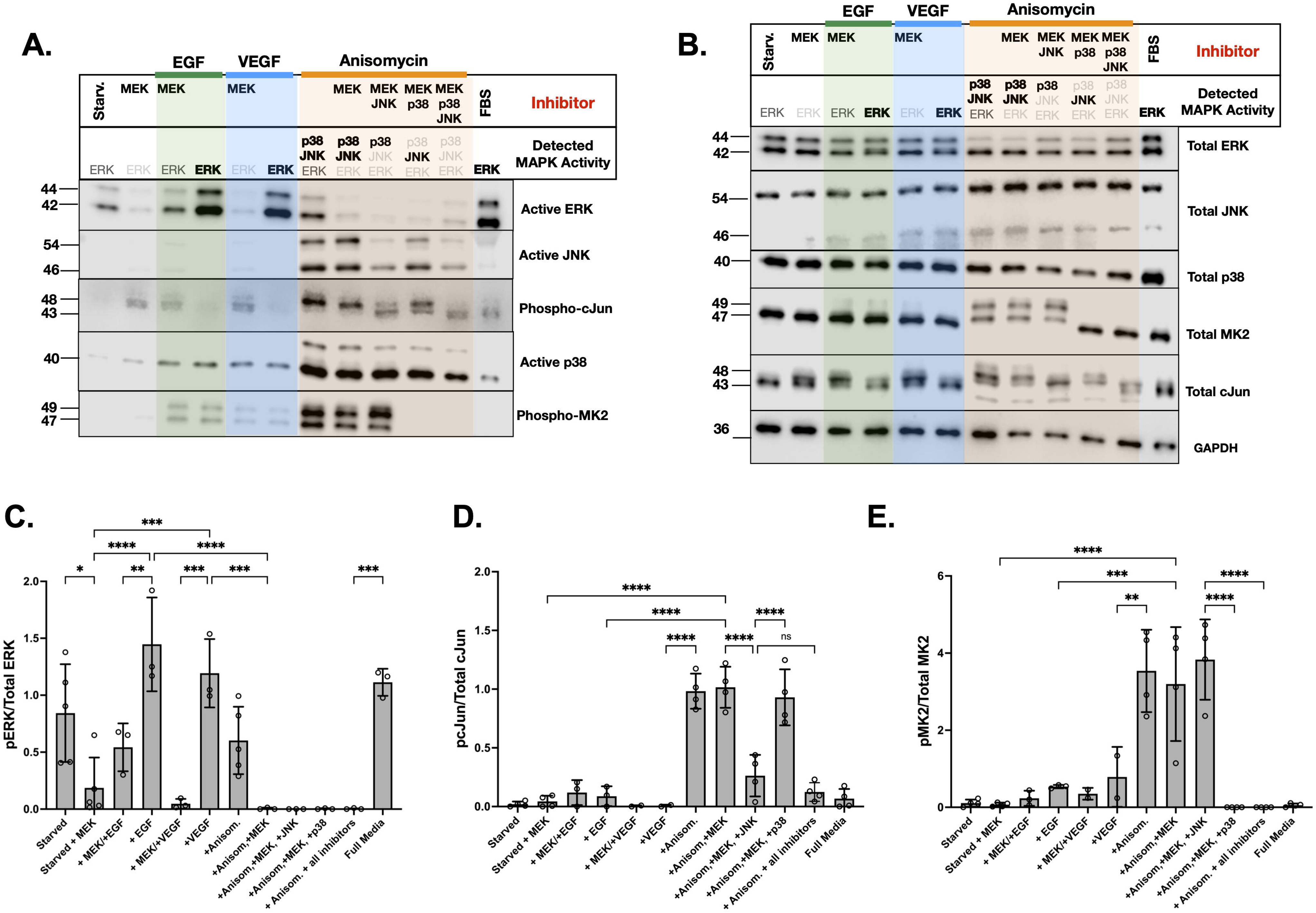
Assessment of MAPK activities in PAEC lysates. All conditions were the same as figure 6, with the addition of treatments with VEGF (blue highlight). **A.** One representative treatment repeat: Western blots of active ERK, JNK, p38, and phosphorylated downstream targets of JNK (c-Jun) and p38 (MK2) in MDCK lysates **B.** Western blots of total MAPKs, c-Jun, MK-2, and the α-tubulin loading control in PAEC lysates used in A. **C.** Quantification of active ERK vs total ERK levels from western blots in A and B. **D.** Quantification of JNK activity through detection of phosphorylated vs. total cJun blots from A and B. **E.** Quantification of p38 activity through detection of phosphorylated vs. total MK-2 from A and B. Statistically significant differences in C-E were determined by ordinary one-way ANOVA: p < 0.05(*), p < 0.005 (**), p < 0.0005 (***), and p < 0.0001 (****). Error bars are shown in SD (standard deviation), from at least three independent cell treatments.

### Assessment of Purified Cx43CT Phosphorylation by MAPKs from MDCKs

To test phosphorylation of full length Cx43 in MDCKs, we initially attempted to transfect WT and serine mutants into this cell line (see materials and methods). Unfortunately, upon cell treatments to activate individual MAPKs, levels of ectopically expressed Cx43 varied significantly as MAPK phosphorylation is known to target Cx43 GJs for degradation. In addition, we struggled with high non-specific signals when using all four phosho-Cx43 antibodies (pS255, pS279, and pS282 - data not shown). To overcome these issues, we utilized a hybrid kinase assay approach, using purified WT GST-Cx43CT protein immobilized on the glutathione resin, and treated the resin with MDCK lysates containing activated ERK, p38 and JNK (**Fig. 8A**). Lysates of MDCKs treated with various activation/inhibition conditions (**Fig. 6**) provided robust and specific levels of Cx43 phosphorylation at S279 and S282 (**Fig. 8A**, quantitated in **8B**). Unfortunately, the signals for S255 and S262 phosphorylation were yet again largely nonspecific (data not shown). Nevertheless, this hybrid approach allowed us to delineate the effects of the ERK signaling from the contribution of the p38 or JNK pathways. Upon activation of ERK, high levels of both S279 and S282 phosphorylation were detected (**Fig. 8A and B**). When both the p38 and JNK signaling pathways were activated and basal levels of ERK activity were eliminated by treatment with the MEK1/2 inhibitor (**Fig. 6**), strong levels of S279 phosphorylation continued to persist, while phosphorylation of S282 dropped to basal level (**Fig. 8**). When p38 was inhibited (and JNK was the only active MAPK), S279 remained phosphorylated. However, when JNK activity was inhibited and p38 was the predominant kinase, S279 and S282 phosphorylation returned to basal levels. Moreover, we were able to identify that under full serum conditions, due to more basal ERK activity, detectable levels of S279 and S282 phosphorylation were present (**Fig. 8A-B**). Taken together, these results indicate that ERK phosphorylates both S279 and S282, JNK prefers S279, and the contribution of p38 is minimal in this cell line.

**Figure 8:**
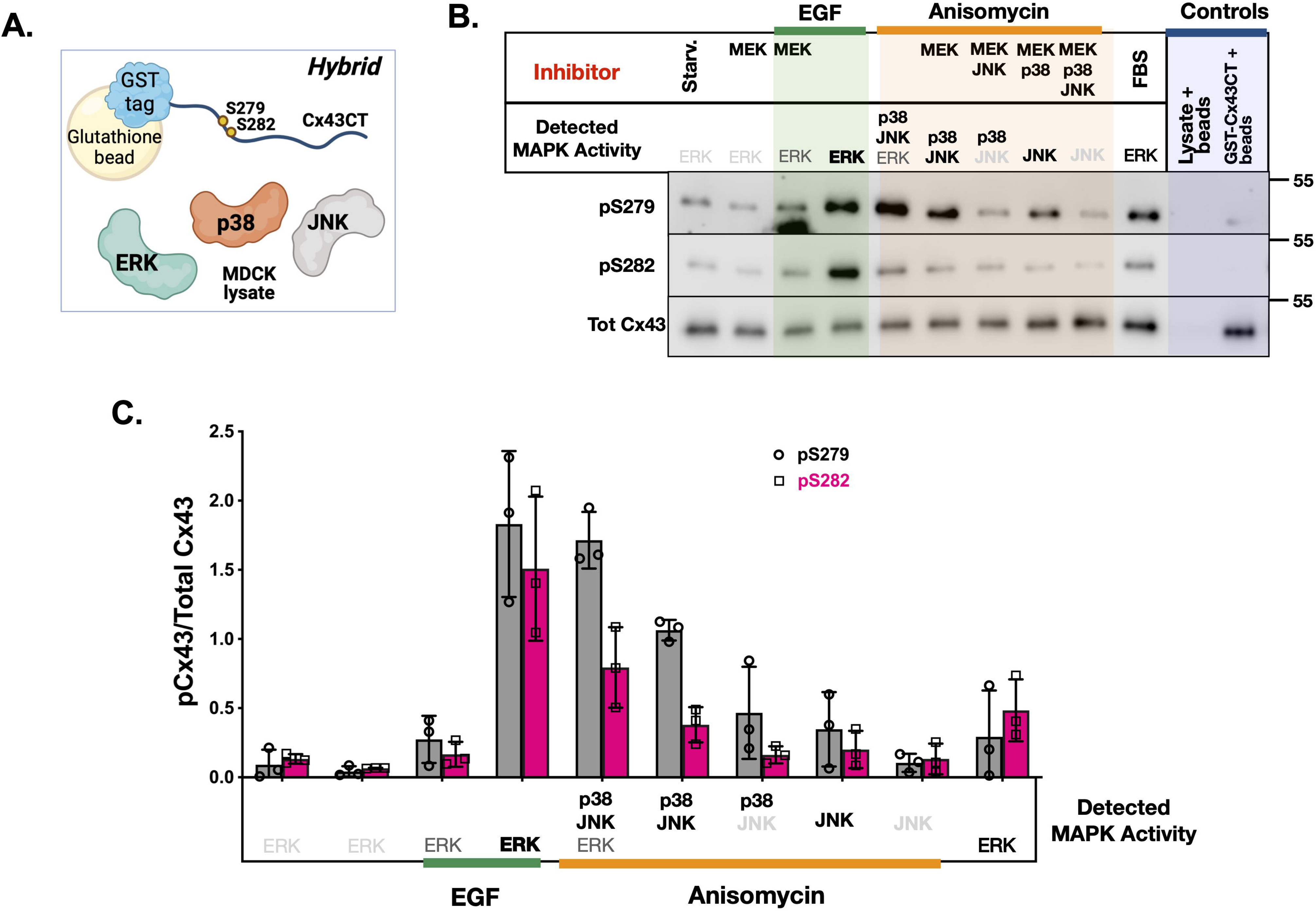
Hybrid kinase assays between purified and immobilized GST-Cx43CT proteins and MAPKs from MDCK lysates. **A.** Cartoon representation of the hybrid approach in assessing phosphorylation of the immobilized, GST-Cx43CT (on glutathione beads) by activated MAPKs from mammalian cell lysates. **B.** A representative hybrid experiment. Levels of total and phosphorylated (pS279, and pS282) Cx43 proteins on glutathione beads post-treatment with activated or inhibited MAPKs. EGF treatments are marked in green, anisomycin treatments are in orange. The presence of inhibitors is indicated. ‘Starv.’ – treatment with lysate from cells treated with FBS-free starvation media; ‘FBS’ – treatment with lysate from cells grown in full media. Negative controls are in blue: cell lysate on glutathione beads and pure GST-Cx43WT on beads. Detected MAPK activity for each condition is summarized using darker, bolded font (higher activity) vs. lighter, thinner font to indicate lower/basal activity. **C.** Quantification of pS279 and pS282 signal from (A.) normalized to the total Cx43 signal. All conditions are in the same order as in (A.) Data are from three independent cell treatments and hybrid kinase assays (n=3). Error bars are in SD (standard deviation). Statistical analyses were not performed due to the use of three different antibodies in WBs. Figure A was generated in BioRender.com.

### Differences in Phosphorylation Preference of Endogenous Cx43 in PAECs

To assess Cx43 phosphorylation by MAPKs in a more native environment, we utilized PAECs, an endothelial cell line that expresses high levels of endogenous Cx43 (**Fig. 9A**). Importantly, all three electrophoretic forms (P0, P1, and P2) have been successfully observed in whole cell lysates by total Cx43 WB in this cell line.^40,48^ Therefore, we hypothesized that the levels of total and phosphorylated Cx43 could be more easily detected in PAEC as compared to our attempts to ectopically express Cx43 in MDCKs . In addition, like in MDCKs, we were able to establish clear activation and inhibition conditions for each MAPK (**Fig. 7**).

**Figure 9:**
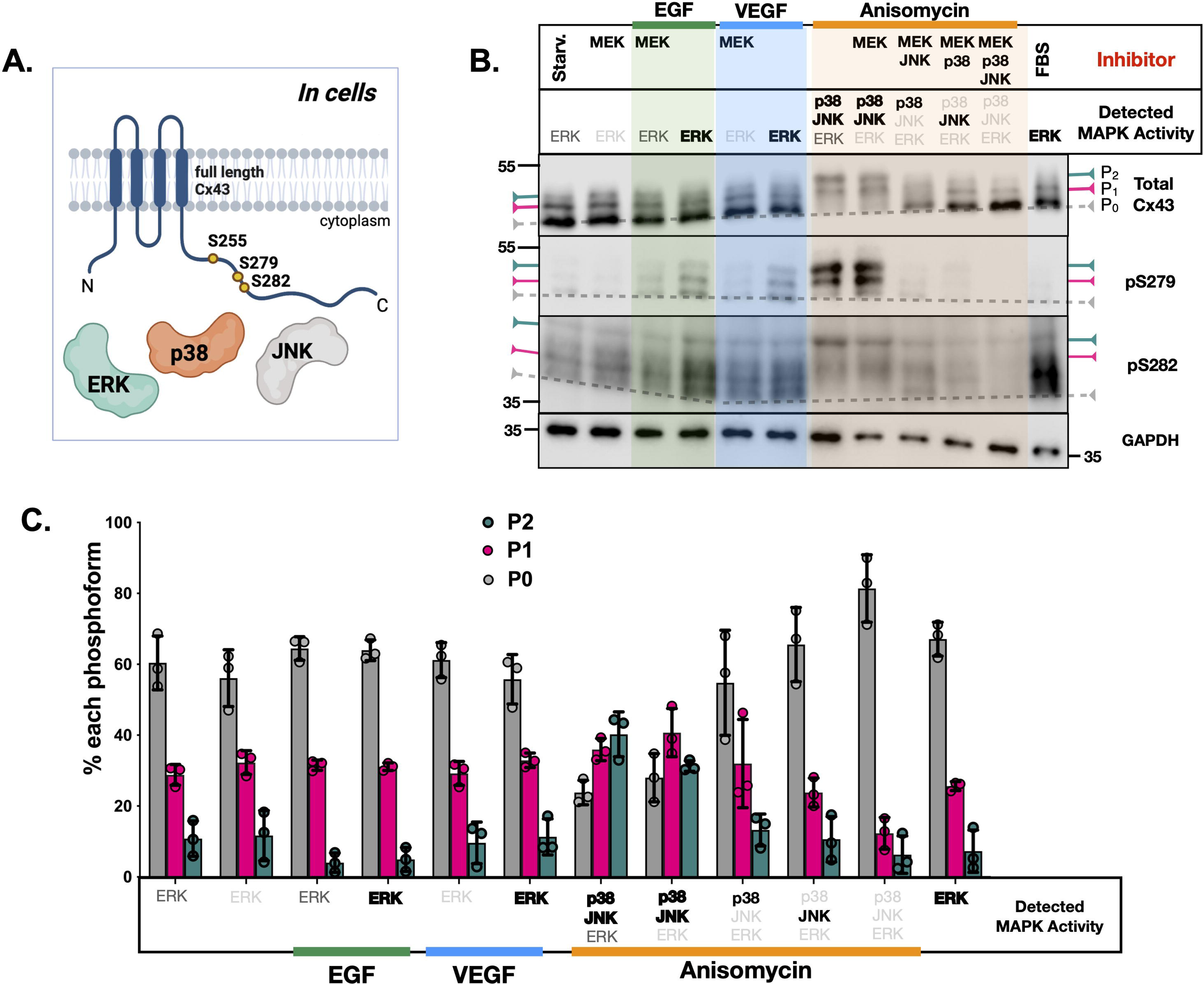
Phosphorylation of endogenous Cx43 in PAECs. A. Cartoon representation of the approach to studying full-length Cx43 phosphorylation by activated MAPKs. (representative experiment). **A.** Detection of phosphorylated (pS279 and pS282) and total Cx43 in PAEC lysates upon activation/inhibition of MAPKs. EGF treatments are marked in green, VEGF - in blue, anisomycin - in orange. The presence of inhibitors is indicated. ‘Starv.’ – lysates from cells treated with FBS-free starvation media; ‘FBS’ –lysate from cells grown in full media. Electrophoretic forms (P0, P1, and P2) are indicated. Detected MAPK activity for each condition is summarized using darker, bolded font (higher activity) vs. lighted, thinner font to indicate lower/basal activity. **B.** Quantification of the total Cx43 signal and the individual electrophoretic forms (P0, P1, and P2) within. Data are an average of three independent cellular experiments (n=3). Error bars are in SD (standard deviation). Two-way ANOVA statistical analyses with all p-values were not included for clarity but are available in Table S2. Figure A was generated in BioRender.

Across 12 treatment conditions, we observed strong signals for phosphorylation at S279 and S282, while also separating the three phosphorylation isoforms in all three western blots (pS279, pS282, and total Cx43) (**Fig. 9B**, quantitated in **9C**). Phosphorylation on S279 was detectable when ERK was activated with EGF or with VEGF, while the highest phosphorylation at this site was observed when both JNK and p38 were active. Inhibition of either p38 or JNK eliminated pS279 phosphorylation almost entirely, while inhibition of both abrogated it fully. (**Fig. 9B**). On the other hand, phosphorylation at S282 was the highest under EGF, VEGF, and especially under full media conditions (FBS). pS282 phosphorylation was present when both JNK and p38 were active and was higher under ‘p38 alone’ than ‘JNK alone’. Taken together, these results demonstrate that ERK can phosphorylate both S279 and S282; p38 can also phosphorylate both sites but to a smaller extent. Importantly, JNK demonstrates the lowest ability to phosphorylate S282 in this cell line (**Fig. 9B**). It is also evident that under full media conditions, pS282 is heavily phosphorylated, with some of the contribution likely coming from a kinase other than ERK (pS282 blot: FBS vs. EGF or VEGF, **Fig. 9B**).

Upon assessment of the mobility shifts of the total Cx43 protein, we observed that activation of ERK failed to produce significant shifts to P2 form, with the bulk of the Cx43 signal in P0 and some in P1 (**Fig. 9B** – total Cx43 WB, quantitated in **Fig. 9C**). However, when both JNK and p38 become activated, Cx43 signal is found mostly in P1 and P2. The addition of either JNK or p38 inhibitor causes an increase in P0 and a decrease in P1 and P2. Once both JNK and p38 are inhibited, levels of P1 and P2 return to basal levels, collapsing largely back into P0 (Total Cx43 WB, **Fig. 9B**, quantitated in **9C, p-values reported in Table S2**). Cx43 phosphorylated at S279 migrates mostly in the P1 and P2 forms (especially under anisomycin treatments when both JNK and p38 are active), while phosphorylation at S282 is observed in all three forms under active ERK conditions and in P1 and P2, when JNK and p38 are active (**Fig. B**, pS279 vs. pS282 WB). Unfortunately, we encountered the same issues with non-specific signals when probing for S255 and S262 phosphorylation as we did in MDCKs (data not shown). Despite these limitations, our results in PAECs indicate that JNK and p38 are the major contributors to electrophoretic mobility shifts to P1 and P2, and to the highest level of pS279 phosphorylation.

### Effects of MAPK Activity on Cx43 Gap Junction Function

To assess if MAPK-phosphorylated Cx43 gap junctions exhibit any changes in channel activity, we carried out scrape-loading Lucifer yellow (LY) dye transfer assays in PAECs across all MAPK activating and inhibiting conditions that influence Cx43 phosphorylation (**Fig 10)**. Appropriately treated PAECs (as described in **Fig. 7**) were scraped in the presence of 0.1% LY and the distance of the LY spread away from the cut within 10 minutes of injury was measured (**Fig. 10A**). MDCKs - because they do not express any Cx43 - were used as a negative control; the LY dye in these cells did not move beyond a single cell layer, indicating lack of cell-cell communication (**Fig. 10B**). Dye transfer of starved cells without any inhibitors was higher than in the presence of the MEK inhibitor (210 vs. 250 µm), indicating that removal of any basal ERK activity significantly increases Cx43 channel opening (Starved only vs. +MEK_in_, **Fig. 10C**, quantitated in **10D**).

**Figure 10:**
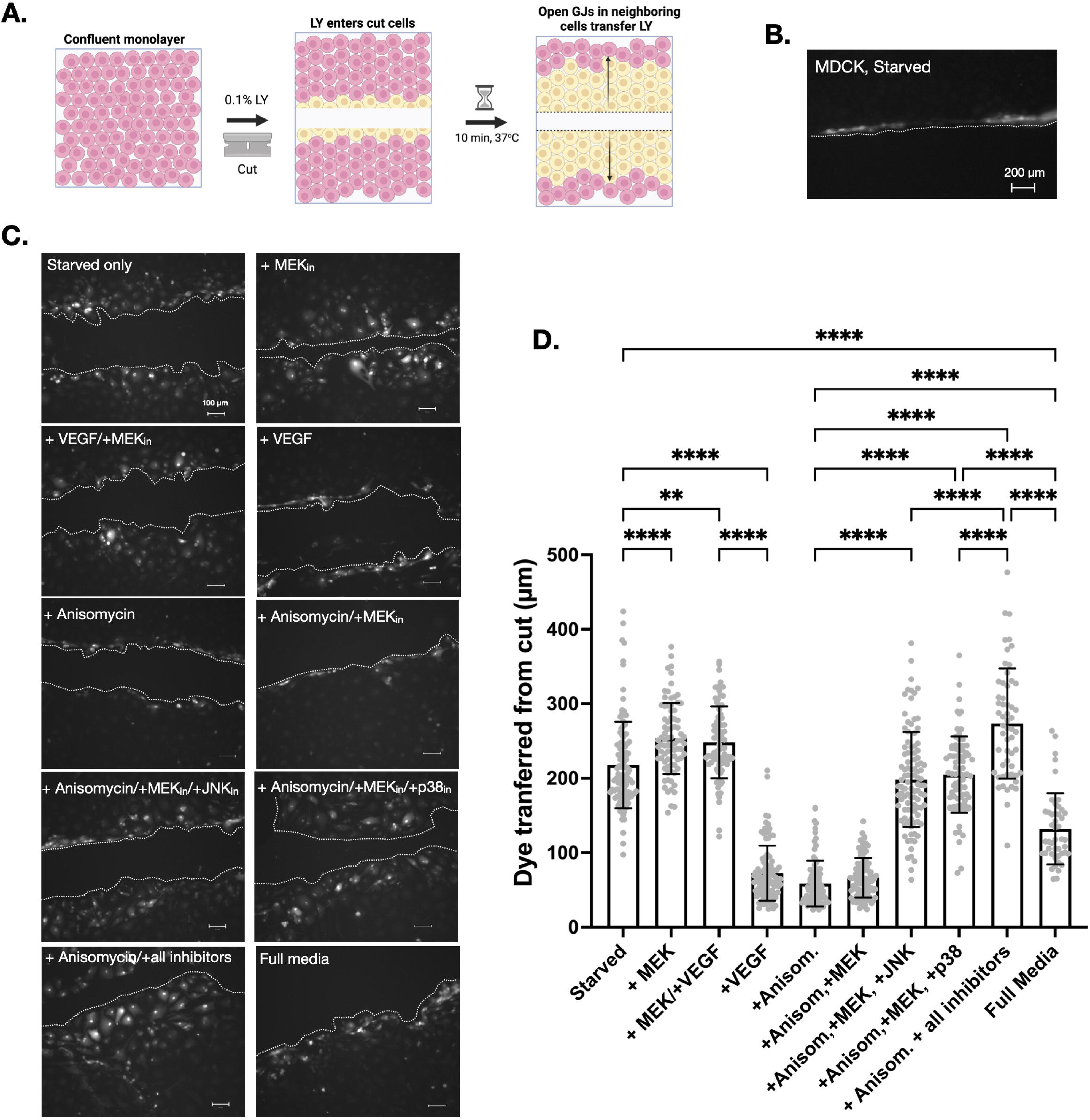
Lucifer yellow scrape loading dye transfer assays. A. Dye transfer principle. B. Negative control – MDCK cells do not have any endogenous Cx43, and thus, cannot transfer dye. The dotted line indicates the edge of the cut. C. A representative biological replicate of PAECs that were treated to inhibit or activate MAPKs (as described in Fig. 8) and scraped in the presence of 0.2 % Lucifer yellow (LY) dye. The samples were imaged on an inverted fluorescent microscope using a 20x objective. Scale bar: 100 µm. **D.** The quantification of three independent biological replicates. Error bars are reported in standard deviation (SD). Statistically significant differences were determined by ordinary one-way ANOVA: p < 0.05(*), p < 0.005 (**), p < 0.0005 (***), and p < 0.0001 (****). Error bars are shown in SD (standard deviation), from at least three independent cell treatments. Figure A was generated in BioRender.

Upon treatment of cells with VEGF (that triggers ERK activation) or treatment with Anisomycin (that triggers JNK and p38 activation), dye transfer dropped to below 80 µm (∼ 1-2 cell widths), indicating GJ closure. GJs remained closed in the presence of anisomycin, even when MEK_in_ was included (Anisomycin + MEK_in_). The addition of either JNK inhibitor or p38 inhibitor increased dye transfer to what was observed with starved cells (200-210µm). However, the addition of both p38 and JNK inhibitors (Anisomycin +MEK_in_ +p38_in_ +JNK_in_) caused a significant increase in dye transfer ability, indicating that when all three MAPKs are inhibited and Cx43 is not phosphorylated, GJs remain open and functional. The same level of cell-cell communication was observed under Starved + MEK_in_ condition, where none of the MAPKs were active). Cells grown in full media exhibited dye transfer distances that were statistically higher than under active MAPK conditions (+VEGF, +Anisomycin), but statistically lower than any condition where only one MAPK was active (for example: starved, where basal ERK activity was present or when JNK/MEK or p38/MEK inhibitors were used) - **Fig. 10C**, quantitated in **10D**. Taken together, these functional assays indicate that activation of ERK, JNK, or p38 has a significant impact on GJ channel closure, driven by phosphorylation of Cx43 by these MAPKs.

## DISCUSSION

Phosphorylation of Cx43 (specifically by ERK) at S255, S262, S279, and S282 is known to mediate GJIC by controlling GJ channel closure, Cx43 ubiquitination, and interaction with endocytosis machinery. However, the individual contributions of each serine phosphorylation by all three MAPK family members have been difficult to pinpoint.^8,10–13,20,22,23,39^ Our study provides a thorough and systematic assessment of substrate specificity by each MAPK toward Cx43, utilizing progressively complex approaches: all purified components (**Figs. 2-4**), in a hybrid mixture of purified and cellular proteins (**Fig. 8**), and directly in mammalian cells (**Fig. 9**). A combination of three complementary techniques (electrophoretic mobility shifts, WB analyses with phospho-specific Cx43 antibodies, and LC-MS/MS phosphoproteomics) together with the use of alanine and glutamic acid mutants help simplify data interpretation of more complex cellular experiments (**Figs. 6-9**). Our work also establishes that T290 - a previously observed (in published mass spectrometry data sets)^49–51^ and predicted^52,53^ phosphorylation site can be phosphorylated by ERK. (**Figs. 2D**, **S6** and **Table S1**). Lastly, we link Cx43 phosphorylation by each member of the MAPK family to GJ closure and downregulation of GJIC (**Fig. 10**).

While we were not surprised that ERK’s specificity toward Cx43 was very different from JNK and p38, significant differences observed between the two stress-activated kinases indicate that JNK and p38 may each be able to regulate Cx43 function in its own way. Kinase assays with purified proteins demonstrated ERK’s ability to phosphorylate all five sites. Purified JNK strongly preferred S255, S262, S279 (with low phosphorylation of S282 and T290), while p38 phosphorylated S262, S279, and 282 (but had a low specificity toward S255 and T290) – **Fig. 2B-D**). Our use of two cell lines allowed us to home in on the cell-line-specific differences; in MDCKs Cx43 phosphorylation is highly driven by ERK (**Fig. 8B-C**), while in PAECs, Cx43 phosphorylation is more dominated through the actions of JNK and p38 (**Fig. 9B-C**). Nevertheless, under active JNK conditions – particularly in the MDCKs - JNK yet again preferred S279 over S282 **(Fig. 8B)** – a result consistent with what we observed in kinase assays with purified components **(Fig. 2)**.

Our experiences with Cx43 mutants as controls (and high background signal in pS255 and pS262) lead us to caution others against the use of Cx43 phosphospecific antibodies (on pure proteins and in more complex mixtures) without additional approaches and alanine site mutants as negative controls. The use of alanine vs. glutamate mutants in gel-shift assays also provided us with evidence that when S255, S279, and S282 are all mutated to alanine, treatment with active ERK does not electrophoretically shift Cx43 to P1/P2, even if S262 is still available for phosphorylation - an observation seen by Warn-*Cramer et. al. 1998* when expressing such a mutant in a mouse Cx43 knock-out (KO) cells treated with EGF.^14^ Our triple E mutant, on the other hand, runs as a P2 form, with or without ERK treatment (**Fig. 4**).

Because p38 has a very low preference for S255 and Cx43 phosphorylation by p38 does not trigger any electrophoretic shift, even when S279 and S282 are becoming phosphorylated (**Fig. 3**), we conclude that S255 is key for a full shift to P2. However, because we did not assess single S279 or S282 mutants, we are unable to tell if only S279, S282 or both need to be phosphorylated along with S255. Indeed, in S255E mutant treated with active ERK, phosphorylation of both pS279 and pS282 is observed in P1 and P2 forms (**Fig. 4A**, pS279 and pS282 blots). These observations were recapitulated using glutamic acid mutants as positive controls: S255E and S279E/S282E were in the P1 form, while the triple E mutant was in the P2 form (**Fig. 4A**, summarized in **4C**). Our results are in full agreement with *Sirnes et. al.* who observed S255 phosphorylation in the P2 isoform of cell lysate samples of IAR20 rat liver epithelial cells exposed to the phorbol ester TPA (12-O-tetradecanoylphorbol-13-acetate) treatments – a condition when ERK is activated.^54^ For S262 phosphorylation, we can conclude that it is largely found in the P0 or P1 form, but not in P2 (**Fig. 4A**, pS262 blot). Moreover, our triple alanine mutant lost all ability to undergo an electrophoretic shift, even if S262 was still available for phosphorylation (**Fig. 4A**), Finally, for T290, T290E mutant remained in P0, similar to the T290E. However, T290E mutant, when phosphorylated on the S sites by ERK, demonstrates a higher shift than T290A (**Fig. 4B**).

Our results with glutamic acid mimetics (**Fig. 4C**) are in some contrast with the electrophoretic mobility of MAPK site mutants described by *Grosely et. al*.^39^ These researchers utilized aspartic acid (D) mutants of some MAPK sites in two different versions of purified CT of Cx43: a soluble GST-Cx43CT (same as in this study) and TM4-Cx43CT (a membrane-anchored version). The study assessed the effects of mimicking phosphorylation on electrophoretic mobility, changes in secondary structure and backbone dynamics of the CT by circular dichroism (CD) and nuclear magnetic resonance (NMR), respectively. In experiments with the fully soluble mutants, GST-Cx43CT S255D/S279D/S282D mutant migrated in the P1 form, while the S255D/S262D and all single mutants remained in the P0 form. The membrane anchored TM4-Cx43CT S255D/S262D mutant was observed in the P1 form, while the TM4-Cx43CT S282D mutant remained in the P0 form. TM4-Cx43CT mutants of other sites were not tested due to difficulties with their expression in *E.coli*.^39^ Even though aspartic and glutamic acid substitutions have both been commonly used to mimic phosphorylation in Cx43,^39,40,55,56^ it is conceivable that the glutamic acid substitution is able to better recapitulate the effects of serine phosphorylation on Cx43.

Our mass spectrometry analyses largely supported the findings of the gel shift assays and western blots, while providing the added benefit of identifying previously uncharacterized sites (**Figure 3D, Figs. S3-S6, Table S1**). One notable shortcoming of our efforts was an inability to distinguish singly phosphorylated S282 and double phosphorylated S279/S282 peptides from the dominant singly phosphorylated S279. This was due in part to the close proximity of these two serines in the Cx43CT amino acid sequence, a challenge encountered in other mass spectrometry studies of Cx43.^57^ Future efforts could attempt to capture these additional phosphorylation events through improved chromatography (longer HPLC gradients, smaller diameter capillary beads), mass spectrometer methods (m/z exclusion lists), or co-elution studies with synthetic heavy-labeled phosphopeptides.

To the best of our knowledge, our work is the first to identify T290 as a MAPK site (**Fig. 3D**, **Fig. S6**, **Table S1**). Even though T290 fully lacks the P-X-S/T-P or S/T-P motifs typically phosphorylated by MAPKs, our mass spectrometry analyses detected T290 phosphorylation in ERK-treated (and to a lesser extent, JNK and p38-treated Cx43CT) (**Fig. 3D**, **Table S1**, **Fig. S6**). T290 phosphorylation has never been reported in any direct peptide sequencing or mass spectrometry analyses of Cx43.^15,57,58^ For example, work by Axelsen *et al* evaluated Cx43 phosphorylation sites by mass spectrometry from Cx43 samples isolated from ischemic vs. control rat hearts; the researchers observed phosphorylation at 255 and 262, while phosphorylation at S279, S282 and T290 was not detected.^57^ Warn-Cramer and colleagues detected S255, S279 and S282 phosphorylation in recombinant Cx43CT treated with recombinant active ERK.^15^ Cooper *et al* observed phosphorylation at S255 *in vitro* when Cx43 was treated with cdc2 and in cells on peptides containing S255, S279, and S282 residues (although these specific sites were not mapped).^58^

Chen and colleagues utilized an informatics server (NetPhosK1.0) to examine phosphorylation motifs in Cx43. While T290 was predicted as a phosphorylation site, it was attributed to PKC by NetPhosK (Chen *et al*).^59^ Even though the server has since been updated (NetPhos 3.1), it is still limited to a set number of kinase motif predictions (ATM, CKI, CKII, CaM-II, DNAPK, EGFR, GSK3, INSR, PKA, PKB, PKC, PKG, RSK, Src, cdc2, cdk5 and p38 MAPK).^60,60^ We assessed Cx43 motifs through the NetPhos 3.1 server that, yet again, predicted PKC and to a lesser extent, cdc2 as kinases for T290 phosphorylation. We also consulted PhosphositePlus^52,53^ for any references to proteomic data sets that have detected T290 phosphorylation. Indeed, phosphorylation at T290 has been observed in a few high-throughput proteomics (HTP) data sets.^49–51^ Importantly, to the best of our knowledge, our findings are the first to attribute T290 phosphorylation specifically to ERK.

Interestingly, a novel T290 mutation was observed in a patient with erythrokeratodermia variabilis et progressiva (EKVP) - a rare congenital skin disorder genetically linked to mutations in connexin genes.^61^ Namely, the study described one patient who had a combination of P283L and T290N mutations, while a second (unrelated) patient had only a single mutation (P283L) (**Fig. 11** - red triangles). While not tested, it is likely that the P283L mutant may not become readily phosphorylated on S282 (as it lacks the canonical phosphorylated motif for S282 phosphorylation - **Fig. 1A** and **B**). Both patients exhibited Cx43 mislocalization (intracellular rather than in the plasma membrane) and the patient with the double mutation demonstrated a more severe EKVP phenotype.^61^ A more recent Cx43 functional study by the Laird laboratory of single (P283L) and double (P283L/T290N) Cx43 mutants expressed in rat epidermal keratinocytes and HeLa cells concluded that these mutations either extend the residency of Cx43 GJs in the plasma membrane or slow down Cx43 internalization and degradation.^62^ These results, together with our identification of T290 as an ERK site, indicate the potential importance of T290 in the regulation of Cx43 internalization and degradation together with the well-characterized effects of S255, S262, S279 and S282 in these cellular processes.^11,12,48^ However, the authors did not test phosphorylation at any serine sites in their P283L mutant (such as S282) and were unable to conclude that a combination of both mutations (P283L/T290N) is the sole reason for yielding a more severe phenotype of EKVP described by Li *et al*.^61,62^

**Figure 11:**
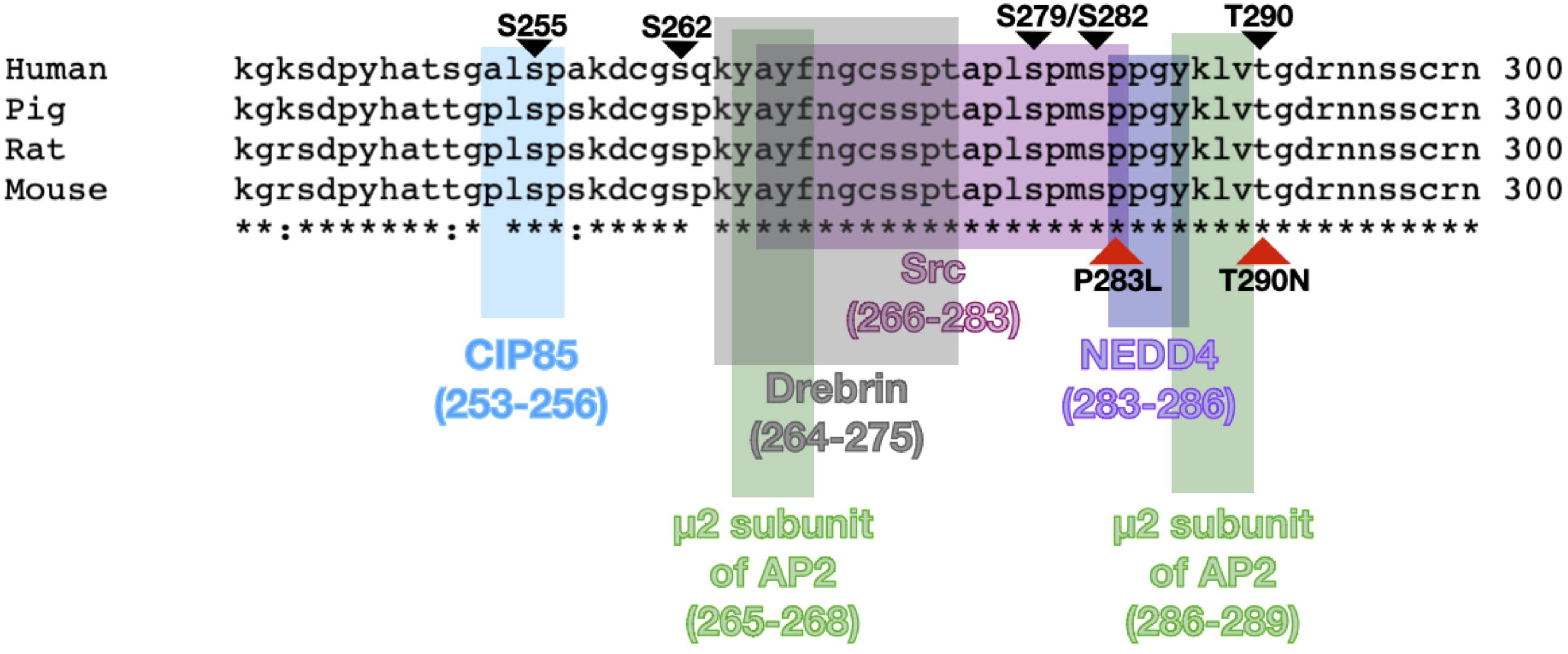
**A.** Sequence alignment (generated using Clustal Omega Multiple Sequence Alignment tool) between human, porcine, rat, and mouse Cx43CT sequence (amino acids 241-300). S255, S262, S279, S282, and T250 are indicated with black triangles. Mutants observed in EKVP patients (P283L and T290N) are in red triangles (Ref. 61). Known binding sites of CIP85, Src, NEDD4, AP2, and Drebrin are highlighted (Ref. 66).

Many excellent *in vitro*, cellular, and *in vivo* studies have demonstrated combinations of Cx43 phosphorylation events under a wide array of activating conditions, such as EGF, TPA (12- O-tetradecanoylphorbol-13-acetate), VEGF (vascular endothelial growth factor) and in many physiological contexts, tissues, and organs.^7,11,12,14,23,34,54,63–65^ What makes the assessment of all this literature difficult is the activation status of all three MAPK family members and whether any sites that were not reported were ever tested. It is thus helpful to have a systematic assessment of phosphorylation events in kinase assays with purified components that help with data interpretation of cellular phosphorylation measurements undertaken under the same treatment conditions (this study) where the activation level of each MAPK is evaluated, and phosphorylation-dead alanine mutants or unphosphorylated pure Cx43CT are used as negative controls.

Finally, this work provides a much-needed confirmation that members of the MAPK family and their signaling pathways are not all created equal when it comes to the phosphorylation of Cx43 on five phosphorylation sites. Importantly, these residues (S255, S262, S279, S282, and T290) lie within or near binding sites of actin interacting proteins (drebrin), trafficking chaperones (CIP85), other kinases (Src), ubiquitin ligases (NEDD4), and clathrin adapter protein 2 (AP2) - summarized in **Fig. 11**.^66^ While we measured levels of open GJs through LY dye transfer studies upon activation of each MAPK (**Fig. 10**), it will be important to further delineate the impacts of each MAPK pathway (ERK, JNK and p38) on all five sites in the fine-tuned regulation of Cx43 protein and GJ function. That includes additional assessments of differences in the Cx43 protein life cycle: its trafficking through the ER and Golgi, oligomerization into hemichannels, GJ formation, key protein-protein interactions (**Fig. 11**), as well as GJ internalization and degradation.

## MATERIALS AND METHODS

### Mutagenesis of Cx43

The pGEX-6p2-WTCx43CT plasmid was a kind gift from Paul Sorgen, University of Nebraska.^39^ Plasmid containing full-length Cx43 for mammalian expression (Cx43-eGFP-NI) was a kind gift from Matthias Falk, Lehigh University. It contains a double-stop codon between Cx43 and eGFP and was used to generate mutants at S255, S279, and S282.^40^ S255A/E, S279/S282A/E, and T290A/E Cx43 mutants were generated using the following mutagenesis primers: S255A: 5’-CAC GCC ACC ACT GGC CCA CTG **<u>GCA</u>** CCA TCA AAA GAC TGC GGA TCT-3’ S255E: 5’-CAC GCC ACC ACT GGC CCA CTG **<u>GAA</u>** CCA TCA AAA GAC TGC GGA TCT-3’ S279A/S282A: 5’ CC TCA CCA ACG GCT CCA CTC **<u>GCA</u>** CCT ATG **<u>GCA</u>** CCT CCT GGG TAC AAG CTG G 3’ S279E/S282E: 5’ CC TCA CCA ACG GCT CCA CTC **<u>GAA</u>** CCT ATG **<u>GAA</u>** CCT CCT GGG TAC AAG CTG G 3’ T290A: 5’ CCT GGG TAC AAG CTG GTT **<u>GCA</u>** GGT GAC AGA AAC AAT TCC 3’ T290E: 5’ CCT GGG TAC AAG CTG GTT **<u>GAA</u>**GGT GAC AGA AAC AAT TCC 3’

To make the triple mutant for bacterial expression, S279/282 mutants were used as templates with the S255 mutagenesis primers. All mutagenesis PCR reactions were amplified using PfuUltra II Fusion HS DNA Polymerase (Agilent). The parental (template) DNA was digested with the restriction enzyme DpnI (New England Biolabs). Plasmid DNA was purified using the QIAquick PCR Purification Kit (Qiagen). The resulting DNA was transformed into DH5𝛼 *E. coli* cells (Invitrogen/Thermofisher). DNA was then isolated using the QIAprep Spin Mini-Prep Kit (Qiagen) and sent for sequencing to ensure that the mutagenesis reactions were successful. Cx43 constructs with a stop codon in the eGFP-NI plasmid were purified with QIAprep Midi Prep Kit (Qiagen) suitable for Lipofectamine2000 (Invitrogen/Thermofisher) transfection following the manufacturer’s transient transfection protocol.

### Protein Expression and Purification of Cx43 WT and Mutants

GST-tagged Cx43CT WT and mutants were expressed in BL21 (DE3) *E.coli* cells. Cx43 expression was induced with 0.75 mM IPTG at an optical density (O.D_600nm_) of 0.6-0.8, and the *E.coli* were grown overnight at 30℃. Bacterial cultures were centrifuged at 8,000 x g for 20 minutes and resuspended in lysis buffer (20 mM Tris-Cl pH 7.5, 150 mM NaCl, 0.5 mM EDTA, 1mM PMSF) for sonication (3 times for 30 seconds at 70% amplitude). The resulting lysate was centrifuged at 12,000 x g to separate the soluble protein fraction and incubated with glutathione resin (GenScript) for 30 minutes while shaking at room temperature (RT). The beads were washed in wash buffer (20 mM Tris-Cl pH 7.5, 150 mM NaCl, 0.5 mM EDTA) and centrifuged at 3000 x g after each wash. The beads were then incubated in elution buffer (20 mM Tris-Cl pH 7.5, 150 mM NaCl, 0.5 mM EDTA, 20 mM glutathione) for 15 minutes at RT. The eluent was collected, dialyzed into a storage buffer (20 mM Tris-Cl, pH 7.5, 150 mM NaCl, 0.5 mM EDTA), and stored at -20°C. Concentrations of GST-tagged Cx43CT proteins were determined by measuring absorbance at 280 nm.

### Protein Expression and Purification of Active MAPKs

Mitogen-activated protein kinase (MAPK) cascades are not naturally found in prokaryotic systems. Therefore, they lack the necessary upstream kinases needed to phosphorylate MAPK such as ERK. The methods for expressing active MAPKs were described previously by Khokhlatchev *et al*. and by Miller *et al*..^67,68^ Plasmids encoding constitutively active, GST-tagged MEK1 (MEK1-pGEX4T1), His-tagged ERK (ERK2-pET28), GST-tagged p38a (p38a-pGEX4T1) and constitutively-active, His-tagged MKK6 (MKK6-pET28) were kind gifts from Benjamin Turk, Yale University, Department of Pharmacology. Plasmids encoding a constitutively active MEKK (MEKK-C-pBR131) and His-tagged JNK2/untagged MEK4 (SAPKa-MKK4-pET15) were a kind gift from Dr. Melanie Cobb, University of Texas Medical Center. Appropriate plasmid combinations were co-transformed into BL21 (DE3) *E. coli* cells using equal amounts of plasmid DNA in presence of both ampicillin (100 µg/mL) and kanamycin (50 µg/mL): ERK2-pET28 with MEK1-pGEX4T1, p38-pGEX4T1 with MKK6-pET28, and SAPKa-MKK4-pET15 with MEKK- C-pBR131.

ERK2 and MEK1 co-expression was induced with 0.4 mM IPTG at OD_600_ of 0.6-0.8, and proteins were expressed overnight at 18℃. Bacterial cultures were centrifuged at 8,000 x g for 20 minutes and resuspended in 10 mL of lysis buffer (20 mM Tris-Cl, 140 mM NaCl, 10 mM imidazole, 10% glycerol, 0.1mM EDTA, 1mM PMSF), and lysed by sonication. The supernatant was incubated with 2 mL of nickel resin (GenScript) at RT for 20 minutes and washed once with lysis buffer and another three times with wash buffer (20 mM Tris-Cl pH8.8, 10 mM imidazole, 140 mM NaCl, 10% glycerol, 0.1mM EDTA). Following the initial washes, the resin was then centrifuged and ERK was eluted in 20 mM Tris-Cl, pH 8.8, 400 mM imidazole, 10% glycerol, 0.1mM EDTA), and dialyzed overnight into dialysis buffer (20 mM Tris-Cl, pH 7.5, 140 mM NaCl, 10% glycerol, 0.5 mM DTT). His-ERK was further purified using 1 mL glutathione resin (GenScript) to remove any residual GST-tagged MKK1.

Purification of active, His-tagged rat JNK2 has been previously described.^69^ GST-tagged p38 expression was induced with 1 mM IPTG at O.D_600nm_ of 0.6-0.8 and the *E.coli* were grown overnight at 30°C. Bacterial cultures (300 mL) were then centrifuged at 8,000 x g for 20 minutes and resuspended in 3 mL of lysis buffer (20 mM Tris-Cl pH 7.5, 10% glycerol, 150 mM NaCl, 1 mM EDTA, 1mM PMSF) and lysed by sonication. Cell lysates were centrifuged at 14,000 x g for 20 minutes to separate the soluble supernatant from the insoluble pellet. To determine the best amount of time that p38 takes to bind to glutathione beads, the supernatants were incubated with Glutathione beads for 20 minutes and 90 minutes at RT or 12 hours at 4°C). After incubation, samples were washed four times in wash buffer (20 mM Tris-Cl, pH 7.5, 150 mM NaCl, 0.5 mM EDTA, 10% glycerol). Following initial washes, 10 mL of elution buffer (20 mM Tris-Cl, pH 7.5, 150 mM NaCl, 0.5 mM EDTA, 10% glycerol, 20 mM glutathione) was added, and samples were rocked at RT for 15 minutes to elute GST-p38. Samples were then dialyzed in storage buffer (20 mM Tris-Cl, pH 7.5, 140 mM NaCl, 10% glycerol, 0.5 mM DTT) and kept at -80°C in 1 mL aliquots. The concentration of all purified proteins was determined by measuring absorbance at 280nm.

### Kinase assays and SDS-PAGE Mobility Shifts

Phosphorylation assays with all pure protein components were carried out at 25°C for 1 hour in 20 mM Tris-Cl, pH 7.6, containing 150 mM NaCl and 10% glycerol, 2 mM ATP and 10 mM MgCl_2_ (termed Kinase Reaction Buffer). GST-Cx43CT WT or S255, S279, and S282 mutant concentrations were kept at 6 µM. The typical reaction volume was 200 µL. Reactions were started by the addition of active MAPKs at a final concentration of 0.5 µM and were quenched with 4x SDS-PAGE Laemmli sample buffer.^70^ Kinase reactions generated for LC-MS/MS analyses were quenched with SDS (0.2% final concentration). Phosphorylation by ERK, p38, and JNK was analyzed via immunoblotting with phospho-specific antibodies against S255, S262, S279, and S282 (see below). For hybrid phosphorylation reactions, 6 µM solutions (500 µL total volume) of pure, GST-Cx43CT (WT or mutants) were added to 200 µL of glutathione beads preequilibrated in 20 mM Tris-Cl, pH 7.6 containing 150 mM NaCl and 10% glycerol (termed Hybrid Assay Buffer). GST-tagged Cx43CT proteins were allowed to bind to the beads for 30 min at RT with gentle rocking. Glutathione beads were washed 3 times in 1 mL of hybrid assay buffer to remove unbound GST-Cx43CT proteins. Lysates (400 µL) of pre-treated MDCK cells (see ‘Lysate Preparation’ section) were then added to GST-Cx43CT proteins and incubated while gently rocking at RT for 1 hour. To ensure that enough ATP and MgCl_2_ were present in these hybrid reaction mixtures, ATP and MgCl_2_ were added to final concentrations of 2 mM and 10 mM, respectively.

Phosphorylation of Cx43 by ERK, p38, and JNK was analyzed by immunoblotting bead samples with phospho-specific antibodies against S255, S262, S279, and S282. The activation status of ERK, p38, and JNK was confirmed by immunodetection of active and total kinase antibodies and of downstream substrates of JNK and p38 in MDCK lysate samples (see below).

### SDS-PAGE and Western Blotting Analyses

Protein samples were resolved on a 4-20% or 10% acrylamide (ThermoFisher) pre-cast Tris-Glycine gels and stained with Coomassie Brilliant Blue stain or transferred for WB analyses onto 0.4 µm nitrocellulose membrane (Bio-Rad) in Tris-Glycine buffer with 20% methanol for 1 hour at 100 V. For the detection of phosphorylated proteins with phospho-specific antibodies, membranes were blocked with 5% BSA in 1x TBS with 0.1% Tween20 (TBS-T) buffer for 1 hour at RT or overnight at 4°C. For the detection of total proteins, membranes were blocked in 5% nonfat milk and TBS-T. Membranes were washed 3x in TBS-T and probed with primary antibodies in TBS-T for 1 hour at RT or overnight at 4°C. After three additional washes, membranes were treated with appropriate goat anti-rabbit-HRP or goat anti-mouse-HRP secondary antibodies (Cell Signaling Technology) at 1:5000 dilution in TBS-T for 30 min at RT. After five additional washes, membranes were treated with ECL Clarity (enhanced chemiluminescence) or ECL Clarity Max Reagent (Bio-Rad) for 5 min and detected on a ChemiDoc^TM^ Gel Imaging System (Bio-Rad) or on the Odyssey XF Imaging System (LiCor). The following primary antibodies were purchased from Cell Signaling Technology and used at 1:2000 dilution: total Cx43, active ERK, total ERK, active JNK, total JNK, active p38, total p38, active MAPKAPK-2 (MK-2) – pThr334 and total MK-2. GAPDH antibodies were purchased from Cell Signaling Technology and used at 1:5000 dilution. ⍺-tubulin antibodies (1:5000), pS262-Cx43 (1:2000), pS279-Cx43 (1:2000), pS282-Cx43 (1:2000) were from ThermoFisher and histidine-tag antibodies (1:2000) were from R&D Systems. pS255 rabbit polyclonal antibodies were a kind gift from Dr. Paul Lampe (Fred Hutchinson Cancer Center, Seattle, WA), and were used at 1:2000 dilution in TBS-T. When probing for purified proteins in kinase assays, a less sensitive ECL reagent was used (Bio-Rad – Clarity ECL), while ECL Clarity Max reagent (Bio-Rad) was used to detect signals from mammalian lysate samples and in the immunoprecipitants.

### Electrophoretic Mobility and Densitometry Analyses

ImageJ (NIH) was used to analyze Coomassie-stained gels for electrophoretic mobility changes and western blots for changes in Cx43 phosphorylation levels at specific serine sites. For mobility shifts, the pixel area between each peak maximum and the higher MW peak minimum was quantified in Keynote (gray in **Fig.3B**). Seven independent kinase assays were quantitated in this manner.

### Kinase Assay Sample Preparation for Mass Spectrometry

SDS was added to kinase assay reactions to a final concentration of 0.2% (v/v). Denatured reactions were then reduced with 10 mM dithiothreitol (DTT) in 50 mM HEPES (pH 8.5) and alkylated with 55 mM iodoacetamide in 50 mM HEPES. Samples were then subjected to single-pot, solid-phase-enhanced sample preparation (sp3) for cleanup and tryptic digestion as follows: samples were incubated with sera-mag speed beads and 50% ethanol for 8 minutes at room temperature, using 10 μg of beads per 1 μg of protein (kinase + substrate). Lysate-bead mix was incubated on a magnetic rack for 2 minutes and the supernatant was discarded. Beads were washed three times with 200 μL of 80% ethanol. Proteins were digested for 18 to 24 hours on bead with sequencing grade trypsin (Promega) in 50 mM HEPES buffer at 1:50 trypsin to protein ratio. Peptides were collected in the supernatant by incubating beads on a magnetic rack, and peptide concentrations were measured by bicinchoninic acid (BCA) assay (Pierce). 5 μg of each sample was lyophilized and labeled with TMTpro isobaric mass tags.

### Phosphoproteomic Analyses of Kinase Reactions

TMT-labeled samples were resuspended in 0.2% trifluoroacetic acid, and pH-adjusted to ≤ 3. Acidified sample was added to Fe-NTA spin columns (ThermoFisher Scientific), and phosphopeptide enrichment was performed as previously described.^71^ Eluates were then subjected to LC-MS/MS on the Q-Exactive Plus as follows: 1) Peptides were separated with a 140 min gradient with 70% acetonitrile in 0.2 M acetic acid. 2) The mass spectrometer was operated with a spray voltage of 2.5 kV. 3) Selected ions were HCD fragmented at normalized collision energy 33%. 4) Full MS1 scans were acquired in the m/z range of 350–2000 at a resolution of 70,000, with maximum injection time of 350 ms. The top 15 most intense precursor ions were selected and isolated with an isolation width of 0.4 m/z. 5) MS/MS acquisition was performed at a resolution of 70,000.

### Phosphoproteomic Data Analysis

Raw mass spectral data files were processed with Proteome Discoverer version 3.0 (ThermoFisher Scientific) and searched against the human SwissProt database and a custom database containing the rat connexin43 C-terminus using Mascot version 2.8 (Matrix Science). Cysteine carbamidomethylation, TMT-labeled lysine, and TMT-labeled peptide N-termini were set as static modifications. Methionine oxidation and phosphorylation of tyrosine, serine, and threonine were searched as dynamic modifications. Peptide spectrum matches (PSMs) for phosphopeptides were filtered for ion score ≥ 15, search engine rank = 1, precursor isolation interference ≤ 35%, and average TMT intensity ≥ 1000. The Proteome Discoverer PhosphoRS node was utilized for improved phosphorylation site localization and assignment, and PSMs were filtered for Isoform Confidence Probability ≥ 0.7. PSMs with missing values were only retained if values were missing for every replicate of a given condition. A TMT intensity of 1000 was imputed for missing values. All remaining PSMs for Cx43 or MAPK family members with a given modification were summed to generate peptide-level TMT intensities. For select Cx43 phosphorylation sites, spectra were manually validated to confirm proper site assignment, and TMT intensity values were obtained directly from the appropriate spectra. Mean-normalized abundances for a given phosphopeptide were calculated by dividing each TMT channel intensity value by the average of these values for the peptide.

**Table S1** has a detailed list of abundances and the amino acid sequence of phosphopeptides from the mass spec run. The raw mass spectrometry data and associated tables have been deposited to the ProteomeXchange Consortium via the PRIDE partner repository with the dataset identifier PXD042575. Data can be accessed for manuscript review with the following credentials:

Username

Password: xmCvof9f

### Mammalian Cell Culture, Cell Treatments, and Lysate Preparation

Porcine Pulmonary Artery Endothelial (pPAEC) cells were a kind gift from Dr. Matthias Falk, Lehigh University. Madin Darby Canine Kidney cells, strain II (MDCK-II)^72^ was purchased from Sigma Aldrich. Both cell lines were cultured in Dulbecco’s Modified Eagle Medium (DMEM) supplemented with 10% fetal bovine serum (FBS), L-glutamine, and penicillin/streptomycin at 37°C and 5% CO_2_. pPAECs express endogenous levels of Cx43, while MDCKII cells do not express any Cx43 but can be transfected with its cDNA.^40,48^ Lipofectamine 2000 (Invitrogen/ThermoFisher) was used for transient, full-length Cx43 WT and mutant transfections into MDCKII cells, following the manufacturer’s protocol. For 30 mm dishes (or 6-well plates), 2.5 µg of plasmid DNA and 5 µL of lipofectamine were used. Human epidermal growth factor (EGF) and human vascular endothelial growth factor - protein 165 (VEGF-165) were purchased from PeproTech/ThermoFisher. EGF stocks (100 µg/mL) stocks were stored in 5% sucrose at - 80°C. VEGF stocks (10 µg/mL) were stored in PBS with 0.1%BSA at -80°C. Anisomycin, JNK inhibitor (SP600125), p38 inhibitor (SB202190), and MEK1/MEK2 inhibitor (U0126) were purchased from Tocris and stored in DMSO (dimethyl sulfoxide) at -20°C. Before any cell treatments, MDCK or pPAEC were starved for 2 hours in FBS-free DMEM media. Inhibitors (SP600125, SB202190, or U0126) were added following starvation and cells were treated for 1 hour at 20 µM of each inhibitor in FBS-free DMEM. JNK and p38 activation was triggered with or without inhibitors for 1 hour using 1 µg/mL anisomycin in FBS-free DMEM. DMSO was used as vehicle control in samples lacking any anisomycin and MAPK inhibitors. ERK was activated with either 20 ng/mL EGF in DMEM (in both MDCKs and PAECs), or with 100 ng/mL of VEGF in DMEM (in PAECs) for 10 minutes. All treatment experiments were carried out at 37°C and 5% CO_2_ and halted by removal of treatment media and the addition of 2x Laemmli SDS sample buffer and boiling for 5 minutes or by rinsing the cells with ice-cold 1x PBS, and lysing them in pre-chilled lysis buffer containing 50 mM Tris-Cl, pH 7.5, 150 mM NaCl, 1 mM EDTA, 0.5% deoxycholate, 1% NP-40, 1 mM β-glycerophosphate, and 1 mM PMSF. Cell culture dishes with lysis buffer were rocked on ice for 25 minutes and scraped. Cell lysates were transferred to pre-chilled 2 mL Eppendorf tubes and centrifuged at 11,000 x g for 5 minutes. The clarified soluble lysate fraction was then used for Cx43 immunoprecipitations or in kinase assays with GST- Cx43CT proteins immobilized on the glutathione beads (as described in the ‘*Kinase Assays’* section above).

### Scrape Loading Lucifer Yellow Dye Transfer Assays

PAECs were seeded on poly-L-lysine coated glass cover slips in 6-well plates and grown to confluence overnight. Cells were then treated to activate or inhibit MAPKs (as described above). Cells were then rinsed in 1x PBS and 0.2% (by wight in 1x PBS) Lucifer yellow (LY – ThermoFisher) was added (1mL/well). Cell monolayers were cut with a razor blade and incubated at 37°C/5%CO_2_ for 10 min. LY was then removed and cells were fixed in 3.7% formaldehyde (in 1x PBS) for 15 min at room temperature. Cover slips were rinsed three times in 1x PBS and dye transfer was imaged on a Nikon Eclipse TE 2000E inverted fluorescent microscope at Lehigh University, Bethlehem, PA (using a 20x objective). Three independent rounds of PAEC seeding, treatments and FY dye transfer assays were carried out, with at least 5-6 fields of view obtained from each repeat. Distances from the cut were measured in ImageJ (NIH) using the ‘measure’ tool and statistically compared. At least 30 measurements were obtained from each biological replicate. A confluent sample of starved MDCKs was treated in the same fashion as a negative control.

### Cx43 Immunoprecipitation from MDCK

MDCK cells transfected with full-length Cx43 cDNA or with Cx43WT-GFP were lysed as described above in 300 µL of Lysis Buffer (50 mM Tris-Cl, pH 7.5, 150 mM NaCl, 1 mM EDTA, 0.5% deoxycholate, 1% NP-40, 1 mM β-glycerophosphate, and 1 mM PMSF) per 3 cm dish. 30 µL of magnetic protein A-beads (BioRad) preincubated with 1 µL total Cx43 antibodies (Cell Signaling Technologies, Cat. # 3512, to immunoprecipitate full-length Cx43) or GFP-trap magnetic agarose (Chromotek, to precipitate GFP-tagged Cx43) were rocked with 200 µL of clarified cell lysate for 1 hour at RT. Beads were washed in 500 µL of lysis buffer three times and eluted in 80 µL of 2X Laemmli SDS sample buffer and boiled for 5 minutes. Samples were then used in WB analyses with phosphoserine or total Cx43 antibodies.

### Statistical analyses

Coomassie gel and western blot signals were quantified using ImageJ software (NIH). Quantifications were carried out on experiments that had at least three independent replicates. GraphPad Prism 9 was used for all bar graphs and statistical analyses. All data are represented as mean ± SD (standard deviation), and significance was tested using one-way or two-way ANOVA, as indicated in the figure captions. All calculated p-values for relevant figures can be found in **Tables S2 and S3**)

## Abbreviations

AP-2: adapter protein 2
Cx43: connexin 43
CDK: cyclin-dependent kinase
cdc: cell-division cycle, gene that encodes CDC1
CK: casein kinase
EGF: epidermal growth factor
ERK: extracellular signal-regulated kinase
GJ: gap junction
GJIC: gap junction intercellular communication
GST: glutathione S-transferase
JNK: c-Jun N-terminal kinase
LY: Lucifer yellow
MAPK: mitogen-activated protein kinase
MDCK: Madin Darby canine kidney
MEK/MKK: MAP kinase kinase
MEKK: MEK kinase/MAP kinase kinase kinase
pPAEC: porcine pulmonary artery endothelial cells
MAPKAPK-2/MK2: mitogen-activated protein kinase-activated protein kinase 2
PKA: protein kinase A
PKC: protein kinase C
TPA: 12-O-tetradecanoylphorbol-13-acetate
VEGF: vascular endothelial growth factor
WB: western blot.

## Acknowledgments

This work was supported through internal funds at Moravian University and by NIH NIGMS R15 (1R15GM148952-01) grant to A.T. The authors are grateful to undergraduate researchers in the Biochemistry II course at Moravian University taught by AT (2017 and 2018) for Cx43CT initial protein purifications and undergraduates at Lafayette College, Easton PA (Molecular Genetics, taught by AT) for preparation of the GST-Cx43CT mutants (2016). The authors are thankful to Lee Graham, Lehigh University Biological Sciences, for the use of the inverted fluorescent microscope for dye transfer assays.

## Supporting information

This article contains supporting information.

## Data availability

All data supporting the results described are in the manuscript and in the supporting information. The raw mass spectrometry data and associated tables have been deposited to the ProteomeXchange Consortium via the PRIDE partner repository (as described in the Materials and Methods).

## Author contributions

**A.T.** - conceptualization of all biochemical and cellular experiments except LC-MS/MS analyses, funding acquisition, project administration, manuscript writing and editing, training/supervision of L.L., L.P., R.T., V.V, S.A., S.S., R.K., H.B., and M.C. - (all undergraduate researchers at Moravian University). **F.W.** - manuscript editing and supervising J.C.-P. **F.W.** and **J. C.-P.** LC-MS/MS conceptualization. **J. C.-P.** - LC-MS/MS data acquisition, manuscript writing, and editing. **L.L.** - cellular and hybrid studies on Cx43 phosphorylation, data analyses, and manuscript editing. **L.P.** - identification of cellular conditions for MAPK activation and inhibition. **R.T.** - kinase assays with all pure MAPKs. **V.V.** - generation and characterization of active p38. **S.A.** - generation and characterization of active JNK. **M.C.** - generation of active ERK and development of the initial ERK/Cx43 assays. **S.S.** – purification of GST-Cx43CT WT and serine mutants. **R.K.** – T290 mutagenesis and purification. **H.B.** – characterization of T290 mutants and assessment of Cx43 phosphorylation in PAECs and MDCKs.

