## supplemental figures for "Differential substrate specificity of ERK, JNK, and p38 MAP kinases toward Connexin 43"

**Supplementary Figures:**

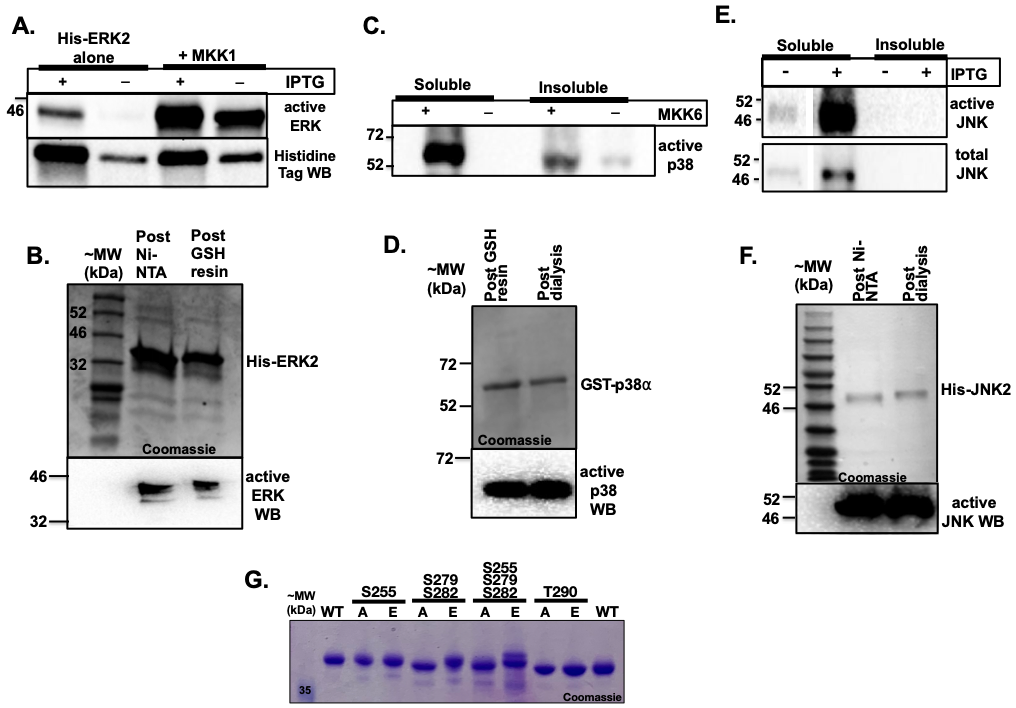

**Figure S1: Expression and purification of active MAPKs and GST-Cx43CT proteins from *E.coli*.** **A.** Levels of activated His-ERK2 when expressed in *E.coli* with or without IPTG induction and with or without coexpression of the upstream MAP2K (MKK/MEK1). Active ERK was detected with antibodies that recognize dual phosphorylated ERK activation loop (pT202 and pY204). Total ERK was detected with antibodies against the poly-histidine tag. **B.** Coomassie-stained gel and active ERK western blot (WB) of purified active His-ERK2 after elution from the Ni-NTA resin and following the cleanup with glutathione resin (to remove residual GST-MEK1). **C.** Expression levels of active and total GST-p38⍺ from *E.coli* with or without coexpresson of the upstream MAP2K (MKK/MEK6). Antibodies against dual phosphorylated p38 activation loop were used (pT180 and pY182). **D.** Coomassie-stained gel and active p38 WB of GST-p38 post purification and dialysis. **E.** Expression of active His-JNK2 with or without IPTG induction. Active JNK was detected with antibodies against dual phosphorylated activation loop (pT183 and pY185). Total His-JNK was detected with total JNK antibodies. **F.** Coomassie-stained gel and active JNK WB of His-JNK2 post purification and dialysis. **G.** Coomassie-stained gel of purified GST-Cx43CT proteins used in this study.

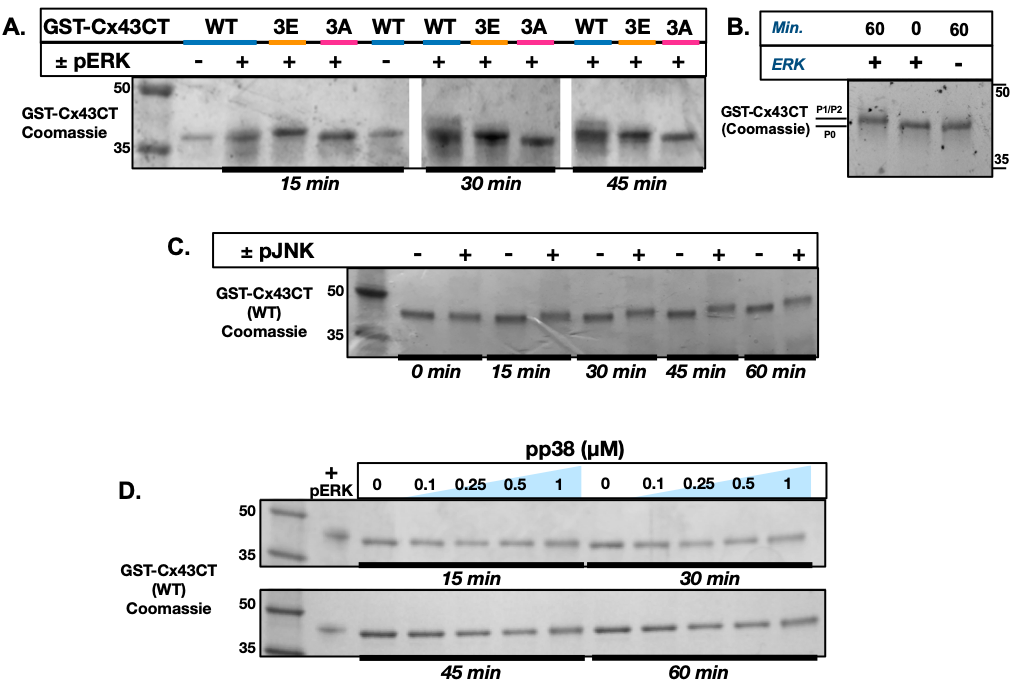

**Figure S2: Initial *in vitro* kinase assays between purified, active MAPKs and GST-Cx43CT** proteins to identify assay conditions (kinase concentration, incubation time) that resulted in complete mobility shifts of Cx43. **A.** Kinase assays between active ERK and GST-Cx43CT (WT, triple A, and triple E mutants – S255/S279/S282). ERK concentration was 0.5 µM, Cx43 concentration was 6 µM. Complete shift of WT to the same mobility as the triple mutant was achieved by 30 min. **B.** Kinase assays between active ERK (0.5 µM) and GST-Cx43CT (WT, 6 µM) after 60 minutes. **C.** GST-Cx43CT WT phosphorylated with active JNK (JNK concentration – 0.5 µM, Cx43 concentration – 6 µM). Robust shift to a higher mobility form was observed after 30 minutes. **D.** GST-Cx43CT WT (6 µM) phosphorylated with active GST-p38. Four time points and four kinase concentrations were tested. Cx43 treated with active ERK was used as a positive control. Despite higher active p38 concentration used (1 µM), no apparent shift in mobility of Cx43 was observed after a full hour, under all concentrations tested. 0.5 µM kinase concentration and 1 hour reaction time were chosen for all subsequent ERK, JNK and p38 kinase assays with pure, GST-Cx43CT proteins (**Figures 2-4**).

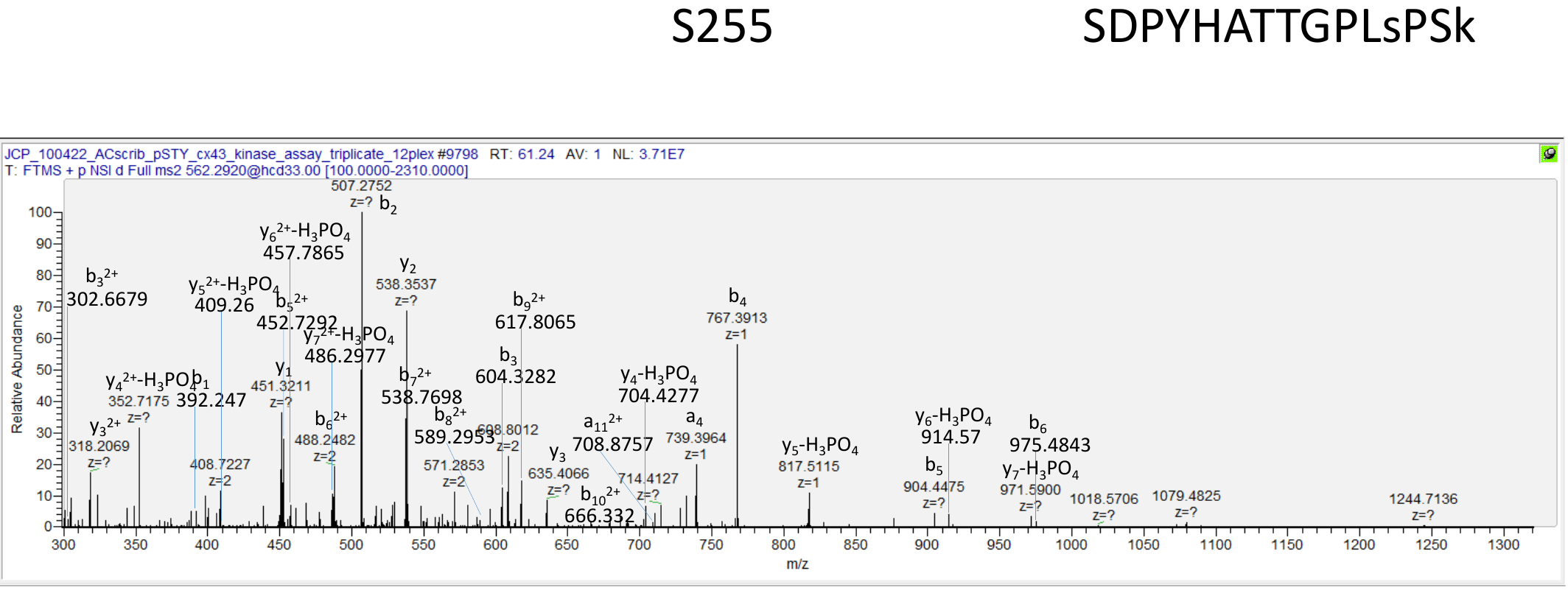

**Figure S3**: **A representative, manually validated MS/MS spectra for S255-phosphorylated GST-Cx43CT.** b‐, y‐, and a- ions and ions corresponding to peptide fragments with neutral losses are annotated. TMT reporter ions from this peptide-spectrum match (PSM) were used to obtain values for relative quantitation. Ions Score: 61, PhosphoRS Best Site Probability- S12(Phospho): 99.99. PhosphoRS Isoform Confidence Probability: 1

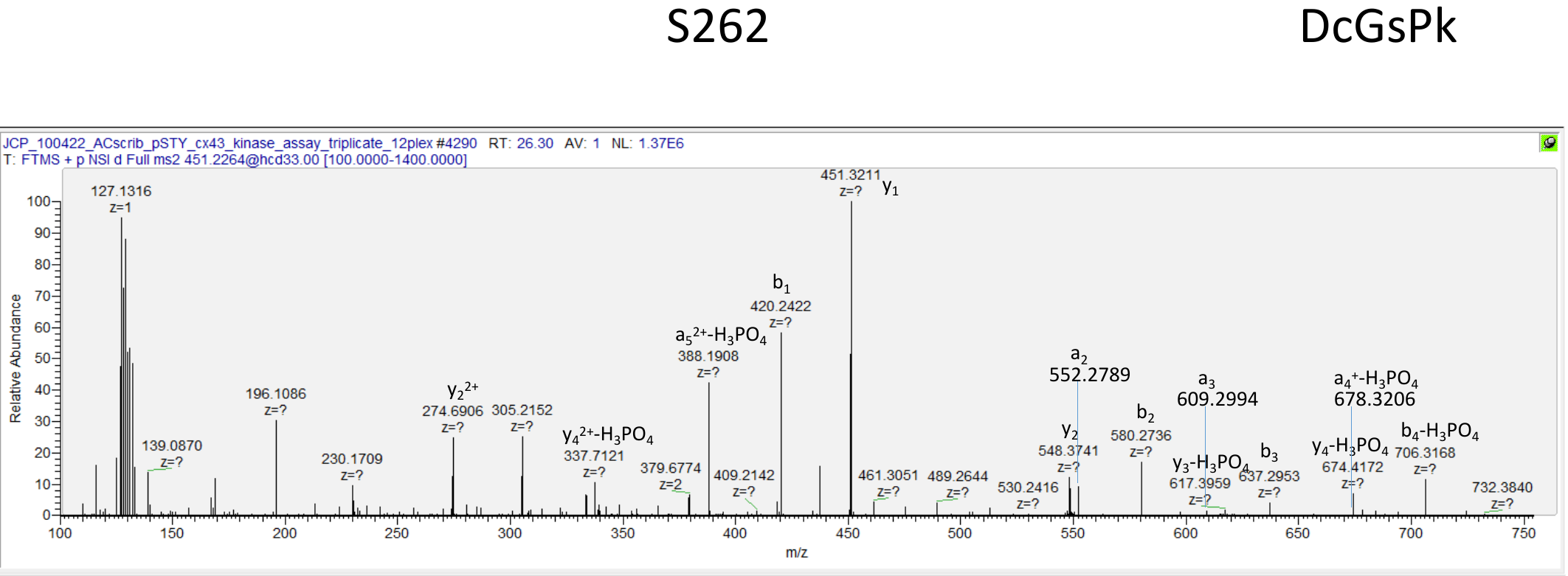

**Figure S4:** **A representative, manually validated MS/MS spectra for S262-phosphorylated GST-Cx43CT**. b‐, y‐, and a- ions and ions corresponding to peptide fragments with neutral losses are annotated. TMT reporter ions from this peptide-spectrum match (PSM) were used to obtain values for relative quantitation. Ions Score: 21, PhosphoRS Best Site Probability-S4 (Phospho): 100. PhosphoRS Isoform Confidence Probability: 1

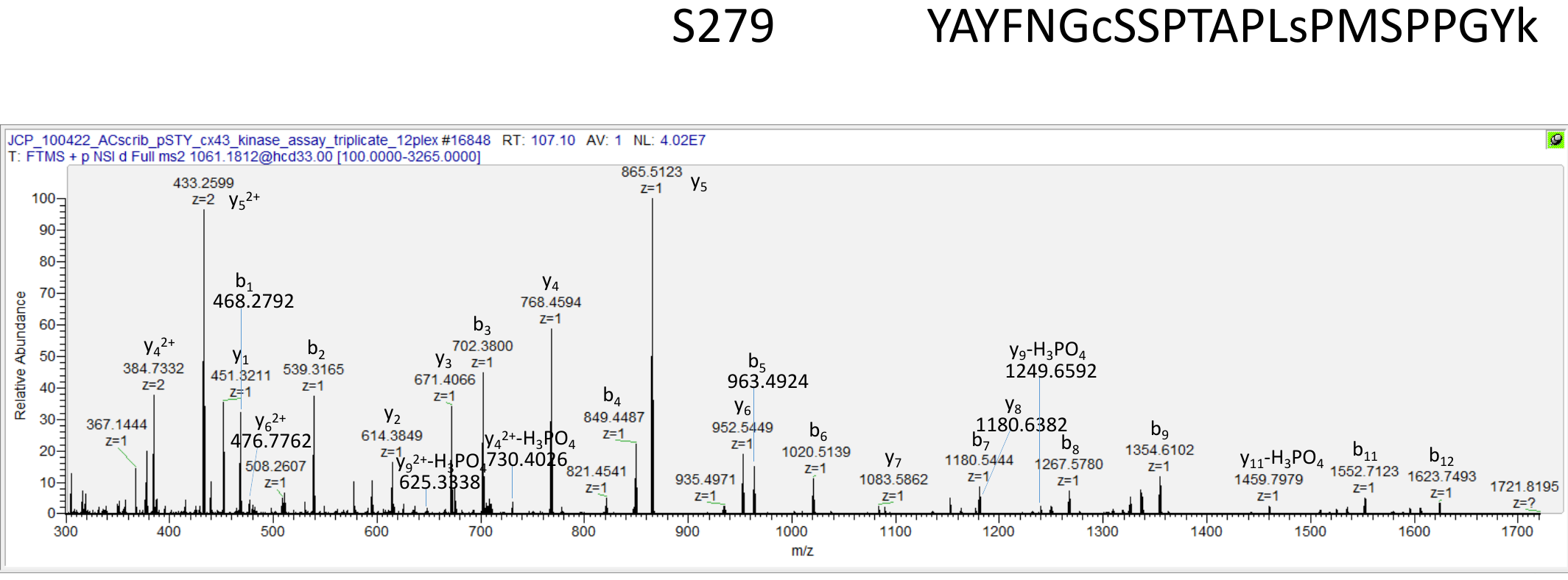

**Figure S5: A representative, manually validated MS/MS spectra for S279-phosphorylated GST-Cx43CT.** b‐ and y‐ ions and ions corresponding to peptide fragments with neutral losses are annotated. TMT reporter ions from this peptide-spectrum match (PSM) were used to obtain values for relative quantitation. Ions Score: 55, PhosphoRS Best Site Probability-S15(Phospho): 100. PhosphoRS Isoform Confidence Probability: 1

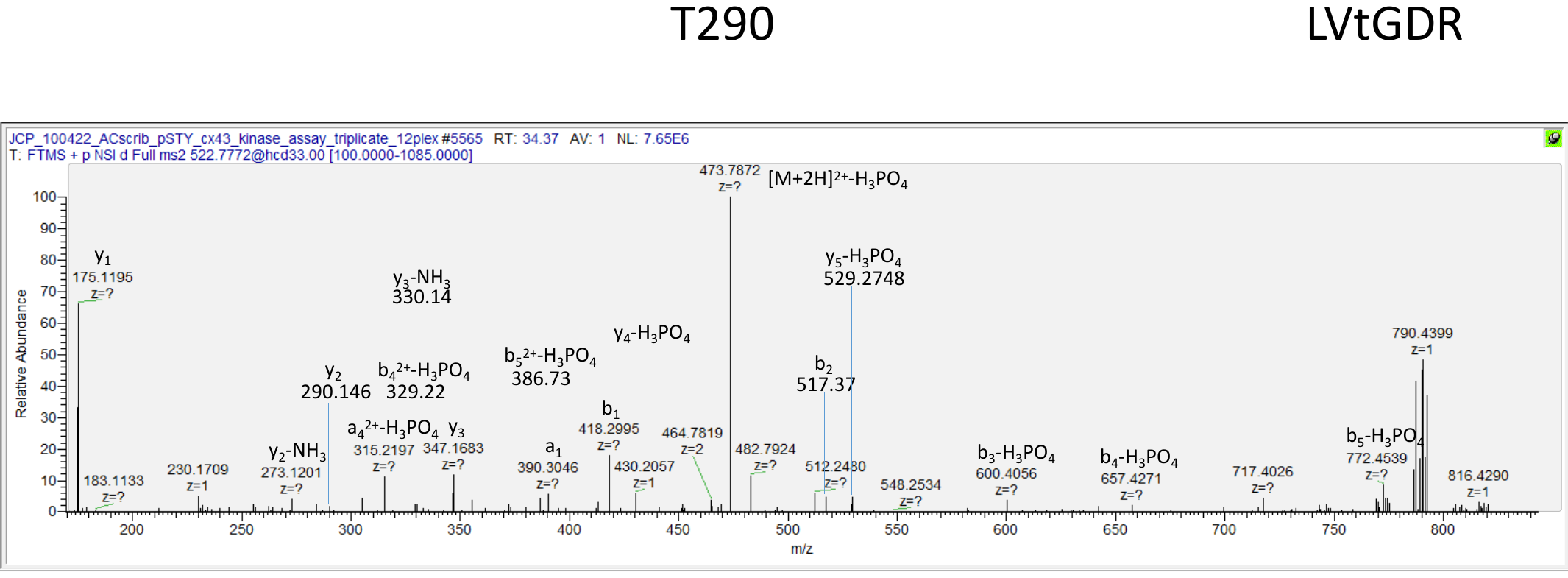

**Figure S6: A representative, manually validated MS/MS spectra for T290-phosphorylated GST-Cx43CT**. b‐, y‐, and a- ions and ions corresponding to peptide fragments with neutral losses are annotated. TMT reporter ions from this peptide-spectrum match (PSM) were used to obtain values for relative quantitation. Ions Score: 12, PhosphoRS Best Site Probability-T3(Phospho): 100. PhosphoRS Isoform Confidence Probability: 1
